# Evolution under pH stress and high population densities leads to increased density-dependent fitness in the protist *Tetrahymena thermophila*

**DOI:** 10.1101/758300

**Authors:** Felix Moerman, Angelina Arquint, Stefanie Merkli, Andreas Wagner, Florian Altermatt, Emanuel A. Fronhofer

## Abstract

Abiotic stress is a major force of selection that organisms are constantly facing. While the evolutionary effects of various stressors have been broadly studied, it is only more recently that the relevance of interactions between evolution and underlying ecological conditions, that is, eco-evolutionary feedbacks, have been highlighted. Here, we experimentally investigated how populations adapt to pH-stress under high population densities. Using the protist species *Tetrahymena thermophila*, we studied how four different genotypes evolved in response to stressfully low pH conditions and high population densities. We found that genotypes underwent evolutionary changes, some shifting up and others shifting down their intrinsic rates of increase (*r*_0_). Overall, evolution at low pH led to the convergence of *r*_0_ and intraspecific competitive ability (*α*) across the four genotypes. Given the strong correlation between *r*_0_ and *α*, we argue that this convergence was a consequence of selection for increased density-dependent fitness at low pH under the experienced high density conditions. Increased density-dependent fitness was either attained through increase in *r*_0_, or decrease of *α*, depending on the genetic background. In conclusion, we show that demography can influence the direction of evolution under abiotic stress.

## Introduction

For many decades, biologists have studied the link between the abiotic environment and the distribution of species on earth, trying to understand why species occur in certain environments and not in others (HilleRisLambers et al., 2012; Dunson and Travis, 1991). Evolutionary biologists more specifically have studied the constraints and potential of species to adapt to their environment and how species respond when changes in their environment occur (Bijlsma and Loeschcke, 2005; Bridle and Vines, 2007). This encompasses research of adaptation to a multitude of abiotic stressors, including salt stress (Gunde-Cimerman et al., 2006; Flowers et al., 2010), heavy metal presence (Shaw, 1994; Klerks and Weis, 1987), thermal stress (Johnston et al., 1990; Angilletta, 2009, chapter 9) and stress associated with drought or the water regime (Kooyers, 2015; Lytle and Poff, 2004). Organisms can respond to such abiotic stress in several ways. They can respond through evolutionary adaptation, by evolving genotypes which match the changed abiotic conditions (Kawecki and Ebert, 2004). They can also adapt through phenotypic plasticity, changing their phenotype to match the abiotic conditions (Westeberhard, 1989). When populations fail to either adapt quickly or to move away — to disperse (Clobert et al., 2001, part 1; Clobert et al., 2012, chapter 1-2) — these populations may be driven to extinction locally. In order to accurately predict local population dynamics and persistence in the context of evolutionary adaptations to abiotic change, it is necessary to understand the speed and direction of evolution in response to changing abiotic conditions, as well as to understand the constraints that such evolution faces.

The question of how populations can adapt through evolution to changing abiotic conditions has a long-standing history in empirical research, both in laboratory experiments as well as field studies (as reviewed in Kawecki and Ebert, 2004). Local adaptation has been recorded in response to different abiotic stressors, across different habitats, and in several taxonomic groups, including plants (Leimu and Fischer, 2008), fish (Fraser et al., 2011), and invertebrates (Sanford and Kelly, 2011). One important environmental impact of human activities is the acidification of natural waters and soils. In the past, acidification has strongly affected natural environments through acid rain (Likens and Bormann, 1974; Likens et al., 1996; Burns et al., 2016). It remains an important abiotic stressor because of the use of fossil fuels and ongoing anthropogenic increase in atmospheric carbon dioxide. Both lead to an acidification of water bodies, oceans in particular, (Caldeira and Wickett, 2003; Raven et al., 2005; Zeebe et al., 2008), with potentially severe consequences for organisms therein. Consequently, recent anthropogenic pressure on the natural environment has triggered increased efforts to understand if and how populations respond to human-induced climate shifts. Reviews of the literature showed that some species evolve to the changing climate, whereas others do not, at least not in the short term (Hoffmann and Sgro, 2011; Franks and Hoffmann, 2012). Ocean acidification has sparked efforts to understand how readily species can evolve to changing pH conditions (Kelly and Hofmann, 2013; Sunday et al., 2014).

Despite a growing body of work, evolution to pH stress is still less well studied experimentally, compared to many other stressors. Evolutionary changes caused by pH shifts have already been studied in the past, and this has typically been done comparatively or through translocation experiments along gradients or between locations differing in pH. For example Derry and Arnott (2007) and Hangartner et al. (2011) showed that copepods and frogs are locally adapted to the pH of their environment. Experimental evolution studies on adaptation to pH stress, although existing, are limited to only few systems that include bacterial model species (Hughes et al., 2007; Zhang et al., 2012; Gallet et al., 2014; Harden et al., 2015) and yeast (Fletcher et al., 2017). For example Gallet et al. (2014) demonstrate how the pH-niche under pH stress evolves through a transient broadening of the niche, followed by specialization. However, many of these studies are focused on adaptation to digestive tracts (Hughes et al., 2007; Harden et al., 2015) or oriented towards industrial application (Fletcher et al., 2017; Zhang et al., 2012). Although controlled experiments can help understand evolutionary adaptation to pH stress, they are still rare (Reusch and Boyd, 2013; Stillman and Paganini, 2015). In addition, existing experiments do not explore important factors that can affect adaptive evolution, such as demography.

Abiotic conditions will alter population performance, and hence also demography. Understanding how demographic conditions influence evolution, specifically the evolution of life-history traits, has led to an extensive body of theory and experiments (Stearns, 1977, 1992). This work has, for example, demonstrated the importance of density-dependent selection and life-history trade-offs between population growth and intraspecific competitive ability (competition-growth trade-offs; Luckinbill, 1978; Mueller and Ayala, 1981; Andrews and Rouse, 1982; Mueller et al., 1991; Joshi et al., 2001). The eco-evolutionary interaction between demographic changes due to abiotic stress, that is, ecological conditions, and adaptation to abiotic conditions, remains less well understood.

Such eco-evolutionary feedbacks highlight that ecological conditions can alter evolutionary trajectories, and, conversely, that evolutionary change can impact ecological conditions (Pelletier et al., 2009; Hendry, 2016; Govaert et al., 2019). Whereas theoretical work has already incorporated the demographic context into evolutionary questions for some time (for a review, see Govaert et al., 2019), empirical work on adaptation to novel conditions still rarely includes the effect of demography on population performance or density explicitly (for some recent examples that do, see Michel et al., 2016; Nørgaard et al., 2019).

In our study, we experimentally explored how four distinct genotypes of the model protist species *Tetrahymena thermophila* evolve when being subjected to either a low pH treatment or a neutral pH treatment (control setting). We explicitly address the question of adaptation to low-pH stress in established populations with densities close to equilibrium. We quantify how evolution changes life-history strategies in four different genetic backgrounds and highlight the importance of trade-offs in life-history traits for understanding how populations adapt to abiotic stress under conditions of high population density, and assess if populations become more similar in life-history strategy.

We can expect directional selection leading to either a maximization of growth rate, or a maximization of competition related traits. When populations experience low competition, the fastest grower likely experiences a selective advantage, and hence we can expect evolution to lead to an increase in the average growth rate. In contrast, when competition is very high due to high population density, strong competitors will likely be under positive selection. Depending on how abiotic stress alters the selection pressures, expected trends in evolution will change. If a stressful abiotic environment affects mostly growth, but does not influence competition, we might expect stronger selection for increased growth. In contrast, if a stressful abiotic environment mostly affects competition (for example, by limiting the amount of available food, or the uptake thereof), we would expect to see stronger selection for investment in competition related traits at lower population densities compared to the optimal abiotic environment.

## Material and methods

### Experiment

#### Study organism

We used the freshwater ciliate *Tetrahymena thermophila* as a model species. Due to its small body size, high population densities and short doubling time of *∼* 4 h (Cassidy-Hanley, 2012; Collins, 2012), *T. thermophila* is well suited for both ecological and evolutionary experiments (e.g. Fjerdingstad et al., 2007; Collins, 2012; Coyne et al., 2012; Altermatt et al., 2015; Jacob et al., 2016). *T. thermophila* is characterized by a high mutation rate in the macronucleus (Brito et al., 2010). This high mutation rate, in combination with large population sizes (here, ranging from *∼* 1 *×* 10^3^ cells/mL to 2 *×* 10^6^ cells/mL), makes the species an ideal model system for adaptation experiments relying on mutation-driven evolution.

We used four clonal genotypes of *T. thermophila* obtained from the Tetrahymena Stock Center at Cornell University. These 4 genotypes are strain B2086.2 (henceforth called genotype 1; Research Resource Identifier TSC SD00709), strain CU427.4 (genotype 2; Research Resource Identifier TSC SD00715), strain CU428.2 (genotype 3; Research Resource Identifier TSC SD00178) and strain SB3539 (genotype 4; Research Resource Identifier TSC SD00660). We selected these strains because they differ strongly in both general life-history strategy and their response to pH stress (see Fig. S2 in Supplementary Information section S2).

We maintained all cultures in axenic SSP medium consisting of proteose peptone, yeast extract and glucose (Cassidy-Hanley, 2012; Altermatt et al., 2015). To avoid bacterial or fungal contamination, we complemented the medium with 10 µg/mL Fungin, 250 µg/mL Penicillin and 250 µg/mL Streptomycin. We added these antibiotics at the start of all bioassays, at the start of the evolution experiment, and at every medium replacement during the evolution experiment (three times per week). At the beginning of the evolution experiment, we cryopreserved the ancestor genotypes in liquid nitrogen and later revived them for bioassays (following the protocol described by Cassidy-Hanley, 2012). Ancestors are from here on referred to as ANC. During the experiment, we maintained cultures at 30 °C, on a shaker rotating at 150 rpm.

#### Evolution experiment

We prepared 32 50 mL Falcon® tubes containing 20 mL of SSP medium with antibiotics. For each of the four genotypes, we inoculated eight tubes with 100 µL of high-density *T. thermophila* culture and let them grow for three days to ensure that populations were well established before starting the evolution experiment. After these three days, we divided the eight replicates of each genotype into two groups, a low pH treatment (from here on abbreviated as LpH) and a neutral pH treatment (hereafter called NpH). At day one of the experiment, we removed 10 mL of culture from all 32 replicate populations and replaced it with 10 mL of SSP medium with antibiotics for the NpH treatment, and with 10 mL of pH-adjusted SSP medium with antibiotics for the LpH treatment. The pH of the pH-adjusted medium used for these 10 mL replacements was prepared by adding 1 M HCl solution to the medium until a pH of 4.5 was reached (1.6 mL of 1 M HCl per 100 mL of SSP medium, for the relationship between added HCl and pH, see Supporting Information section S1). We repeated this regime of medium removal and replacement on every first, third and fifth day of the week for a total of six weeks. Consequently, the pH of the medium for LpH populations was gradually reduced over a period of two weeks, after which it was kept approximately stable at 4.5 for the remainder of the experiment.

#### Genotype revival and common garden conditions

In order to perform all population growth assays of evolved (LpH and NpH) and ancestral (ANC) populations at the same time, we revived the ancestor populations from liquid nitrogen storage. We transferred revived cells to SSP-medium with antibiotics for recovery. We then prepared a common garden treatment. We inoculated common garden cultures for the LpH, NpH and ANC populations (50 mL Falcon® tubes with 20 mL of SSP medium with antibiotics) with 100 µL culture and transferred them to a shaker for 72 h, in order to control for potential plastic or parental effects. This should ensure that any observed phenotypic changes are the result of either de novo mutations, or of highly stable epigenetic effects.

#### Population growth assessment

After culturing all populations in the same environment (common garden), we assessed population growth at low pH (pH 4.5) and neutral pH (pH 6.5) of the assay medium for the ANC (four genotypes, each replicated four times per assay medium pH treatment), and evolved (LpH and NpH) populations (29 surviving populations per assay medium pH treatment) for a total of 90 cultures. We placed these cultures in an incubator, and grew them for seven days. Most populations reached equilibrium density well before the end of these seven days (between 20 and 100 hours after populations started growing; see also section S10 in the Supporting information), which allows us to obtain precise measurements of growth rates and population equilibrium densities.

#### Data collection and video analysis

We sampled populations both during the evolution experiment and during the population growth assessments, to quantify (i) population density during evolution, (ii) intrinsic rates of increase (*r*_0_), and (iii) intraspecific competition coefficients (*α*) for the ANC, LpH and NpH populations. These *r*_0_ and *α* estimates were obtained through fitting of a population growth model, as described below in the section “Population growth model fitting”. During the evolution experiment, we sampled three times per week prior to medium replacement. For the population growth rate assessments of the evolved and ancestral populations, we sampled a total of 10 time-points over a course of the seven days, with more frequent sampling early in the growth phase (four times over two days) to adequately capture the population dynamics. For sampling and analysis, we followed a previously established method of video analysis to extract information on cell density and morphology of our evolved and ancestral populations, using the BEMOVI R package (Pennekamp et al., 2015).

Our population sampling method is adapted from well-established protocols (Fronhofer and Altermatt, 2015; Fronhofer et al., 2017). Briefly, 200 µL of culture was sampled from the population, and if cell density was too high for video analysis, diluted 1/10 or 1/100, because excessive cell density decreases the accuracy of cell recognition during video analysis. We then transferred the culture to a system of microscope slides with fixed capacity, so that a standard volume (34.4 µL) of culture could be measured for all videos. Next, we took a 20 s video at 25 fps (total of 500 frames) using a Leica M165FC stereomicroscope with top-mounted Hamamatsu Orca Flash 4.0 camera. We analyzed our videos using the BEMOVI R package (Pennekamp et al., 2015) to extract the relevant information. Parameters used for video analysis can be found in the Supporting Information (section S3).

### Statistical analyses

All statistical analyses were performed using the R statistical software (version 3.5.1) with the ‘rstan’ (version 2.18.2) and ‘rethinking’ (version 1.5.2) packages (McElreath, 2015).

#### Population growth model fitting

In order to analyze population growth dynamics of ancestral and evolved populations, we fit a continuous-time version of the Beverton-Holt population growth model (Beverton and Holt, 1993). As recently discussed by Fronhofer et al. (2018, see also chapter 5 in Thieme 2003), using this model provides a better fit to microcosm data compared to less mechanistic models (for example an *r*-*K* population growth model, which captures the density-regulation of microcosms less well) and readily allows for a biological interpretation of its parameters. The Beverton-Holt model is given by the equation

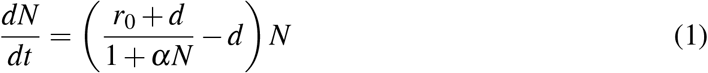

with the intraspecific competitive ability (*α*) being

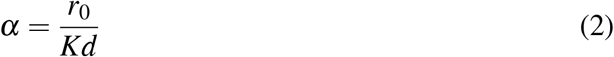

Here, *N* corresponds to population size, *r*_0_ corresponds to the intrinsic rate of increase, *α* to the intraspecific competitive ability (hereafter referred to as competitive ability), and *d* to the death rate of individuals in the population. The *K* parameter in equation (2) represents the equilibrium population density. We adapted Bayesian statistical models from Rosenbaum et al. (2019) to estimate parameter values for *r*_0_, *α*, *d*, and *K* using the rstan package and trajectory matching, that is, assuming pure observation error (see https://zenodo.org/record/2658131 for code). We chose vaguely informative priors, that is, we provided realistic mean estimates, but set standard deviation broad enough to not constrain the model too strongly, for the logarithmically (base *e*) transformed parameters with *ln*(*r*_0_) *∼ normal*(*−*2.3, 1), *ln*(*d*) *∼ normal*(*−*2.3, 1) and *ln*(*K*) *∼ normal*(13.1, 1).

#### Analysis of parameter estimates *r*_0_, *α*, and *K*

In a next step, we analyzed the population growth parameter estimates to determine how our experimental treatments affected them. As intrinsic rates of increase (*r*_0_) integrate birth and death rates and are more reliably estimated than its components (narrower posterior distributions), we here focussed on intrinsic rates of increase and excluded the death rate from further analyses (see also Tab. S10 for summarized posteriors).

To analyse the parameter estimates (*r*_0_, *α*, and *K*), we constructed separate linear models for each genotype, and fit logarithmically (ln) transformed parameters *r*_0_, *α* and *K* as a function of a) the pH of the assay medium, b) general evolution across pH treatments, that is, difference between ANC populations, on the one hand, and evolved populations, on the other hand, c) evolution to specific pH treatments (that is, differences between ANC, LpH and NpH) and d) interactions between pH of the medium and evolutionary changes. This resulted in 16 statistical models for each of the response variables and each of the four genotypes (see Tab. S3 in Supporting Information section S7 for details). Information on priors can be found in the Supporting Information (section S4). Following McElreath (2015, chapter 14), we did not only use our mean parameter estimates, but took their uncertainty into account by modelling both means and errors of the parameters obtained during Beverton-Holt model fitting.

We then compared the models using the deviance information criterion (DIC), a Bayesian implementation of the Akaike information criterion (Gelman et al., 2014) and averaged the posterior predictions of the 16 models based on DIC weights. Next, we calculated the relative importance (RI) of the explanatory variables by summing for each explanatory variable the respective model weights in which this variable is included.

#### Correlation between *r*_0_ and *α*

In order to detect potential correlations between intrinsic rate of increase (*r*_0_) and competitive ability (*α*), we performed a Bayesian correlation analysis using the logarithmically transformed estimates of *r*_0_ and *α* and fitting a multivariate normal distribution. We again used both mean estimates and their errors to account for errors caused by population growth model fitting. To account for plastic effects associated with the pH of the assay medium, we performed the correlation analysis separately for low pH and neutral pH of the assay medium, while pooling the data for all four genotypes and treatments (ANC, LpH, and NpH). Pertinent computer code can be found in the Supporting Information (section S5).

#### Variation in life-history traits

We asked whether evolutionary history altered between-genotype variation in life-history traits (*r*_0_, *α* and *K*) at low and neutral pH of the assay medium. We first calculated for each group (ANC, LpH and NpH) the mean of the natural logarithm of *r*_0_, *α* and *K* over all 4 genotypes, and subsequently calculated the absolute difference between this mean and the observed trait values (*r*_0_, *α* and *K*) of all replicate populations (logarithmically transformed). We then used Bayesian models to calculate whether these differences varied between the treatments (Evolved (general evolutionary change, difference between ANC and all evolved lines), LpH and NpH). To account for potential genotype effects, we also included both models with and without random effects per genotype (random genotype intercepts), leading to a total of 6 models per trait, as shown in Tab. S4 in the Supporting Information section S7. After fitting the models, we compared the models using the Watanabe-Akaike information criterion (WAIC), a generalized form of the Akaike information criterion used for comparison of Bayesian statistical models (Gelman et al., 2014). We then calculated relative parameter importance using WAIC weights.

#### Density-dependent fitness calculation

To assess how the observed convergence in life-history strategy might have arisen, we calculated the population growth rate (*r*) for the LpH and for ANC populations over all observed population densities during the evolution experiment and integrated over these values to calculate a weighted density-dependent fitness estimate. We then used Bayesian models to fit these density-dependent fitness values as a function of a) population origin (ANC or LpH), b) centered intrinsic rate of increase (*r*_0_), and c) an interaction term between *r*_0_ and population origin. Centered *r*_0_ represents the intrinsic rate of increase, rescaled to have its mean at zero, and was calculated by subtracting the mean *r*_0_ from all *r*_0_ values. In this analysis, we also included a random intercept for the different genotypes (details in Tab. S5 in section S7 of the Supporting Information). We fit all five models, starting from the intercept model to the full interaction model. Subsequently, we ranked these models using the WAIC criterion and calculated the relative importance of all explanatory variables based on WAIC weights. The corresponding analysis for the NpH populations can be found in Supporting information section S9.

## Results

We subjected replicate populations of four different genotypes to either low pH (LpH) conditions or neutral pH conditions (NpH), while keeping population densities high over the course of the evolution experiment. Fig. 1 shows the population densities as observed during the experiment. We then tested whether and how evolution changed life-history strategies in all four different genetic backgrounds. Fig. 2 shows the data and model predictions for changes in life-history traits. Next, we tested how life-history traits were correlated and how this may have constrained evolutionary changes. The correlation in life-history traits is depicted in Fig. 3. We then tested for changes in variation of life-history strategy between populations (shown in Fig. 4). Lastly, we tested how evolution of life-history strategies affected density-dependent fitness under the observed densities during the evolution experiment. Fig. 5 shows data and model predictions of density-dependent fitness under low pH conditions.

**Figure 1:**
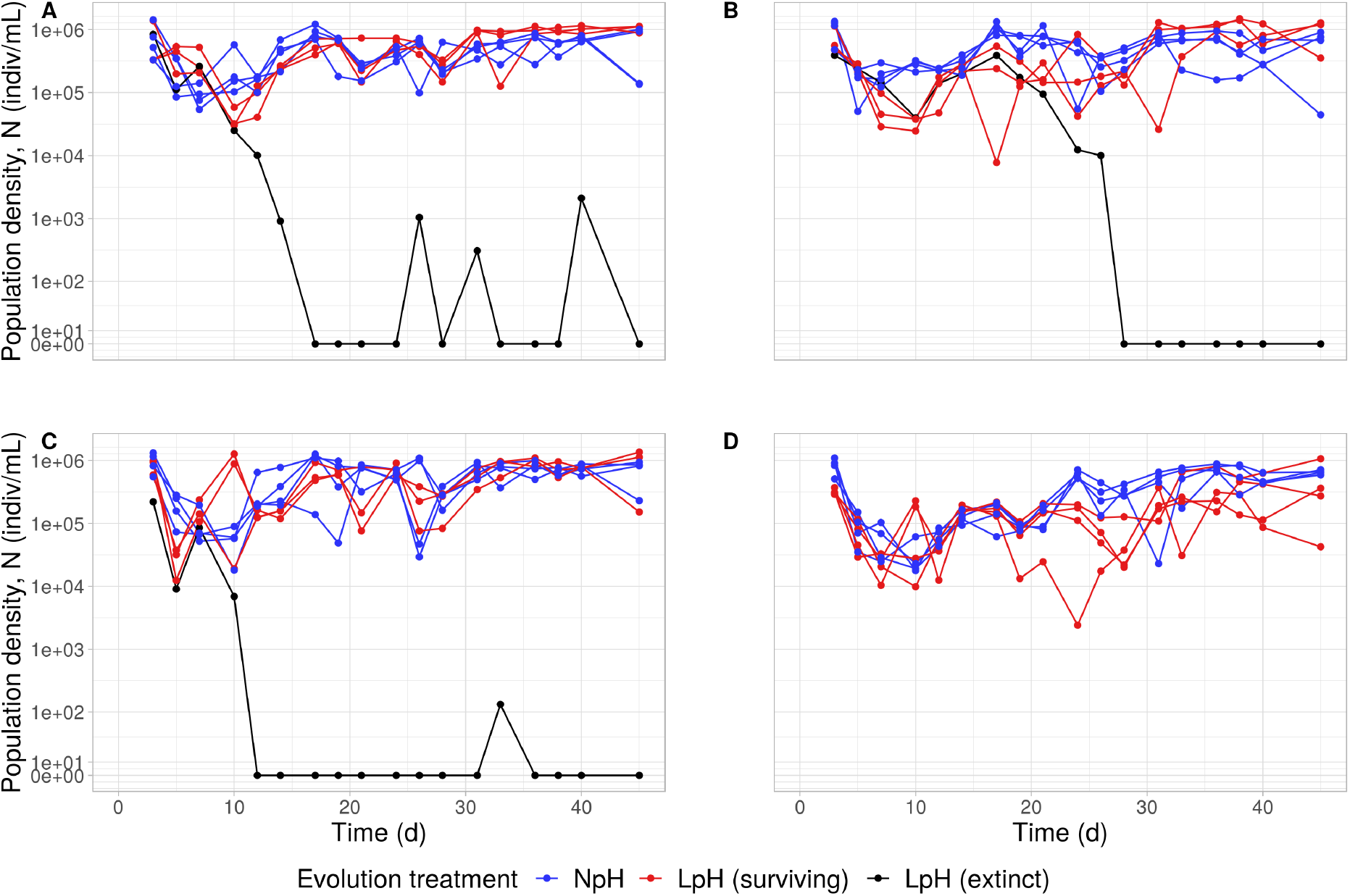
Density dynamics of the replicate populations over the course of the evolution experiment. The y-axis shows the population density (axis is pseudo-logarithmically transformed, to account for 0 values in the dataset), the x-axis the time since the beginning of the experiment in days. Each set of dots connected by a line represents data from a single replicate population. Red and blue symbols correspond to data from populations that survived to the end of the experiment from the LpH (populations evolved under low pH conditions) and NpH (populations evolved under neutral pH conditions) treatments, respectively. Black symbols correspond to data from LpH populations that went extinct. Panel A shows the density dynamics for genotype 1, panel B for genotype 2, panel C for genotype 3 and panel D for genotype 4.

**Figure 2:**
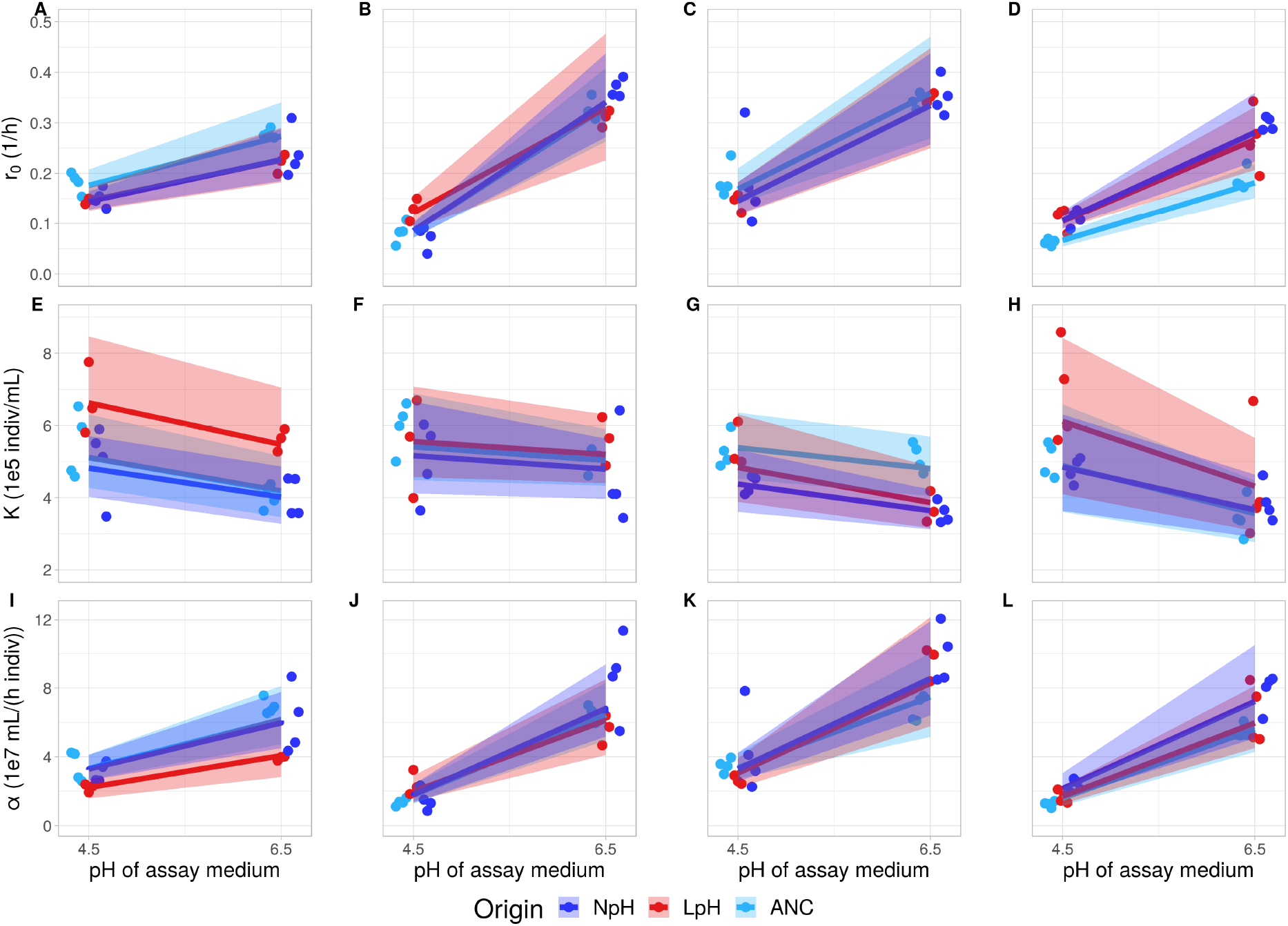
Evolutionary trends in intrinsic rate of increase (*r*_0_; A-D), equilibrium population density (*K*; E-H) and competitive ability (*α*; I-L) for the 4 different genotypes. Each circle represents an estimate of *r*_0_, *K* or *α* (posterior means) from the Beverton-Holt model for one replicate population. Lines and shaded areas represent the averaged posterior model predictions based on DIC weights (means and 95 % probability interval). Light blue = ANC (ancestor populations), dark blue = NpH (populations evolved under neutral pH conditions), red = LpH (populations evolved under low pH conditions).

**Figure 3:**
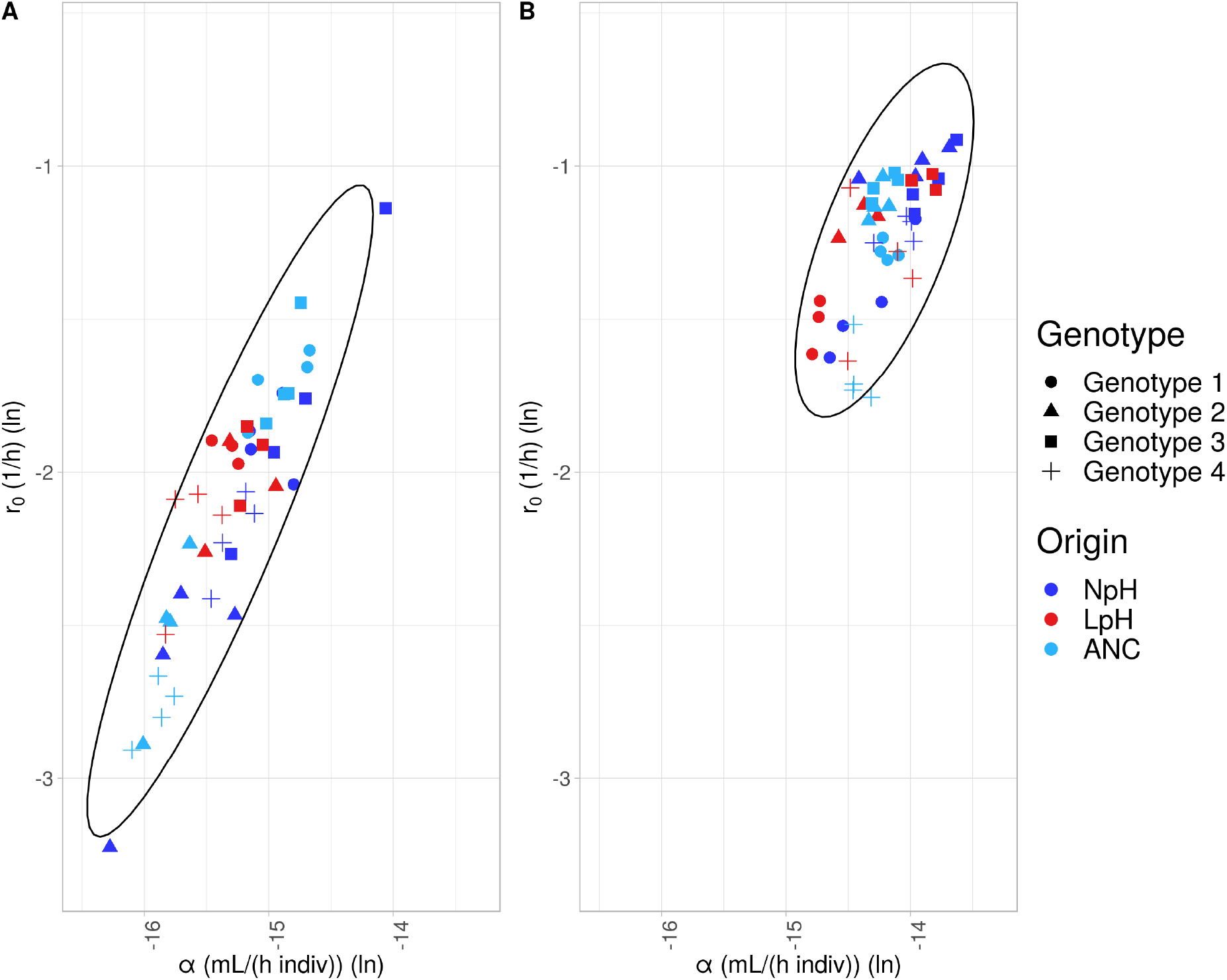
Correlation between the intrinsic rate of increase (*r*_0_) and competitive ability (*α*) at low pH (A), and at neutral pH (B) of the assay medium. Symbols represent the different genotypes (see legend); Light blue = ANC (ancestor populations), dark blue = NpH (populations evolved under neutral pH conditions), red = LpH (populations evolved under low pH conditions). Ellipses represents 95% probability intervals.

**Figure 4:**
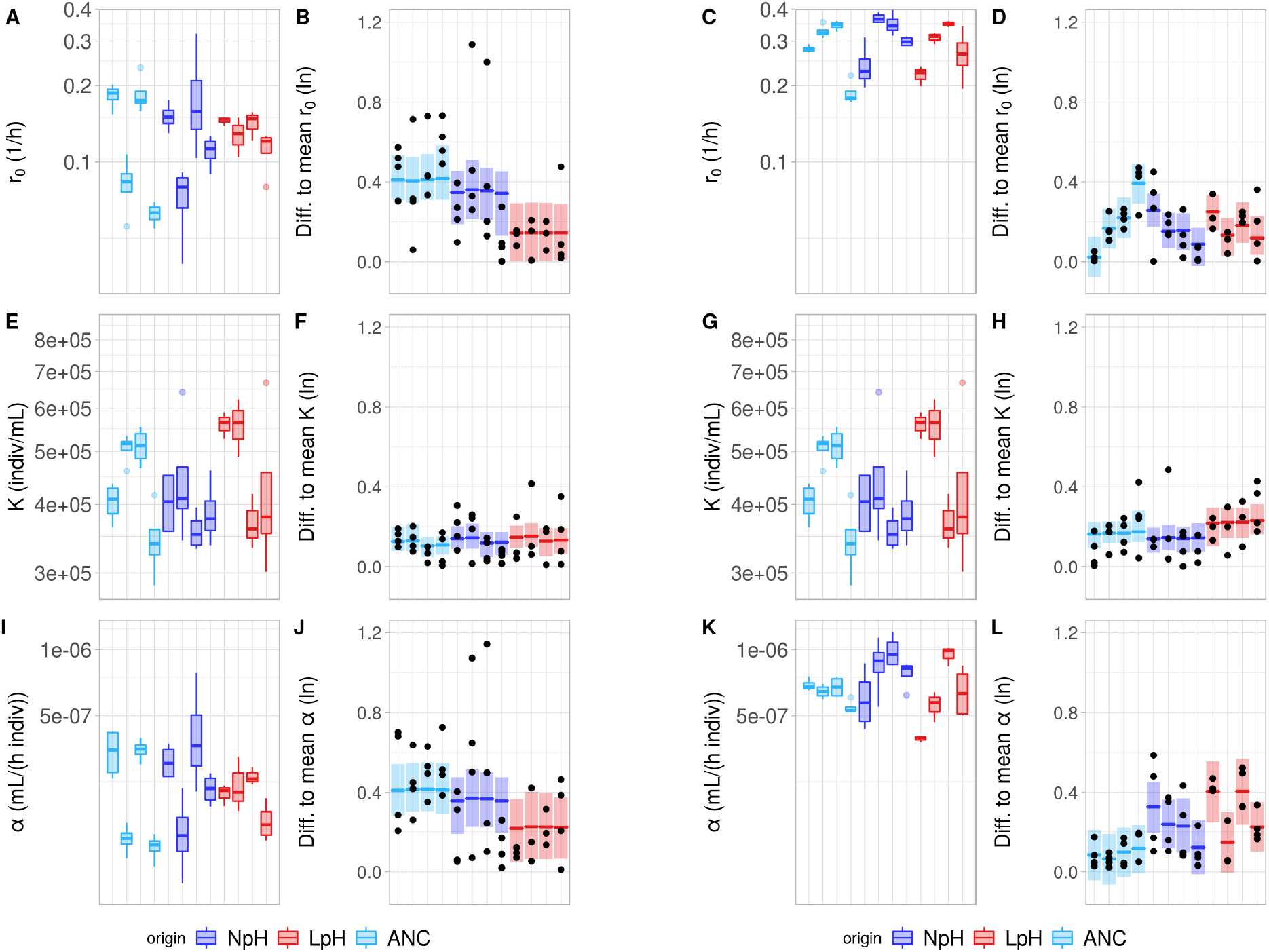
Left half of the figure (panels A,B,E,F,I,J) shows data for growth curves measured at low pH of the assay medium, right half (panels C,D,G,H,K,L) for growth curves measured at neutral pH of the assay medium. Traits shown are intrinsic rate of increase (*r*_0_; A-D), carrying capacity (*K*; E-H) and competitive ability (*α*; I-L). Panels A, C, E, G, I and *K* show *r*_0_, *K* and *α* estimates (1 box plot is 1 genotype). Panels B, D, F, H, J and L show averaged model predictions (mean and 95 % probability interval) of difference between *r*_0_, *K* and *α* estimates and mean per treatment (ANC, LpH or NpH; boxes) and individual datapoints (black dots). Light blue = ANC (ancestor populations), dark blue = NpH (populations evolved under neutral pH conditions), red = LpH (populations evolved under low pH conditions).

**Figure 5:**
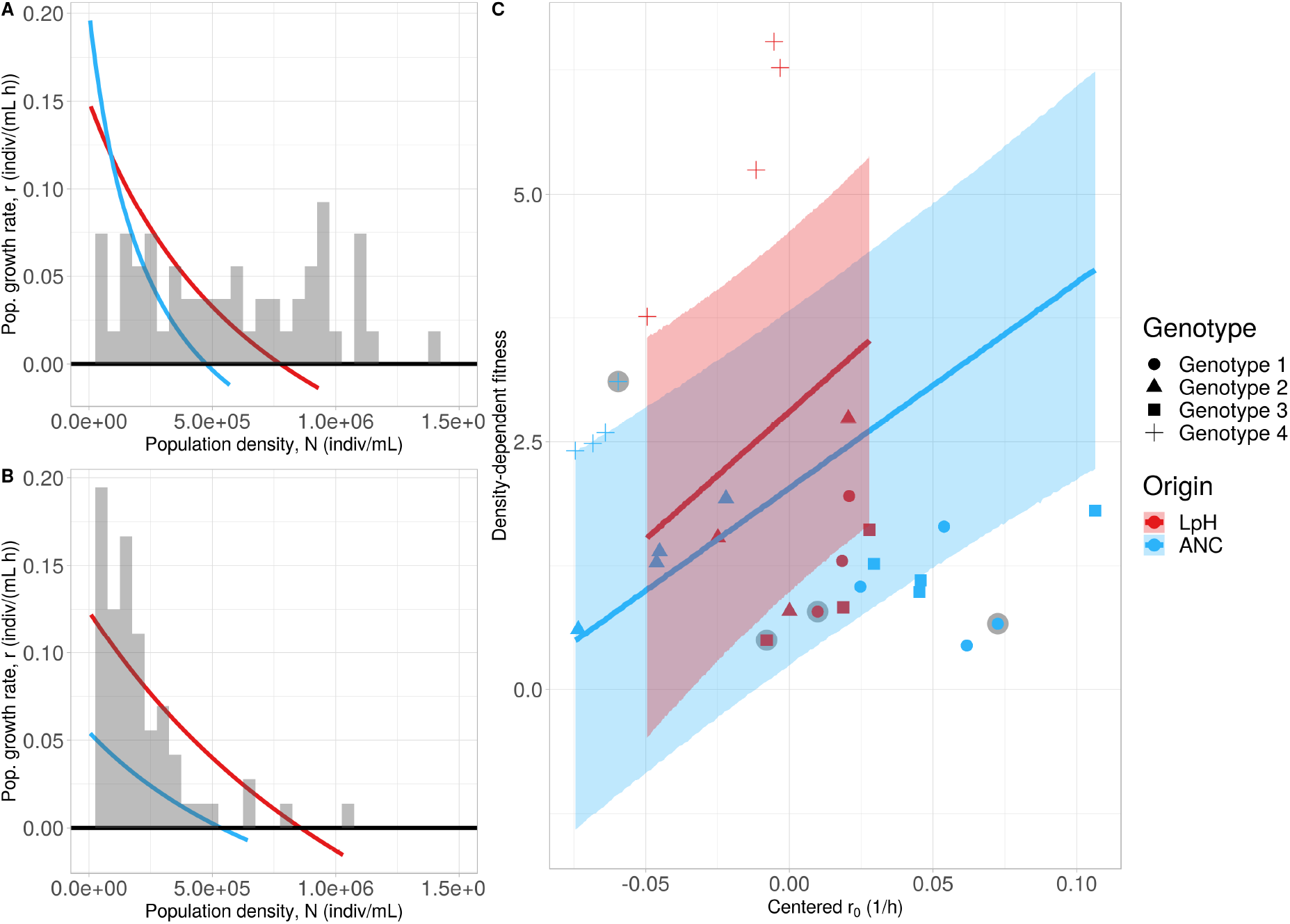
Density regulation functions for selected populations where the intrinsic rate of increase (*r*_0_) evolved to decrease (panel A – genotype 1) or increase (panel B – genotype 4) in LpH populations (populations evolved under low pH conditions). Light blue lines show the density regulation functions for the ANC populations (ancestral populations), red lines for the LpH populations. Grey bars show a histogram of the observed population densities during the evolution experiment for the corresponding genotype. C) Density-dependent fitness depending on the (centered) intrinsic rate of increase (*r*_0_). Symbols correspond to data from LpH (red) or ANC (blue) populations (shape represents genotype, see legend). Symbols surrounded by a grey disc represent the example populations of panels A-B. Lines and shaded areas represent the weighted posterior predictions and the 95% probability intervals for the four genotypes. A visual representation of the density regulation function of all replicate populations can be found in the Supporting Information Fig. S4 in section S6.

### Evolution of life-history traits

During the 42 days of the evolution experiment, population densities ranged from approximately 1 *×* 10^3^ cells/mL to 2 *×* 10^6^ cells/mL (see Fig. 1) and fluctuated around the population equilibrium density due to stochastic variation in death and division rates. Observed densities varied strongly depending on treatment and genetic background. Out of 32 evolving populations, three went extinct during the experiment, all in the low pH treatment (one population each for genotype 1, 2 and 3).

After the experimental evolution phase, we found that all four genotypes showed strong plastic effects associated with the pH of the assay medium (see also Tab. S6 in the Supporting Information section S8). Low pH of the assay medium consistently decreased intrinsic rate of increase (*r*_0_), led to lower competitive ability (*α*), and, as a consequence of this decrease in *α*, to increased equilibrium population densities (*K*) as shown in Fig. 2. This effect of low pH was especially pronounced for *r*_0_ and *α*, where the relative importance values associated with pH of the medium were typically close to one for all four genotypes (see also Tab. S6 in the Supporting Information section S8). The effect of low pH was less pronounced for the equilibrium population density (*K*), specifically for genotype 2.

We additionally found signatures of evolutionary change. These were less consistent than the plastic effects, that is, they differed between the genotypes. Evolution led to an increase in *r*_0_ for genotypes 2 and 4 (Fig. 2B,D). However, for genotype 2 this increase only occurred in the LpH populations. For genotype 4 we mostly observed a general change in all evolving populations and only to a lesser degree specific changes in the LpH and NpH treatments.

LpH led to increased equilibrium population density (*K*) for genotype 1 and genotype 4 (Fig. 2E, H), and a decreased equilibrium population density (*K*) for genotype 3 (Fig. 2G). As equilibrium density is an emergent trait, the changes in *K* were driven both by the changes in *r*_0_ described above and by changes in *α*. Evolution led to lower competitive ability (*α*) for LpH genotype 1 populations (Fig. 2I), to increased competitive ability (*α*) for evolved genotype 2 populations (Fig. 2J), to no clear change for genotype 3 (Fig. 2K), and to increased competitive ability (*α*) for evolved and especially NpH for genotype 4 populations (Fig. 2L). Overall, we detected evolutionary changes in all traits (*r*_0_, *α* and *K*), although direction and strength of change strongly differed between genotypes.

### Variation and covariation in *r*_0_ and *α*

The intrinsic rate of increase (*r*_0_) and competitive ability (*α*) were positively correlated both at low pH and neutral pH of the assay medium (Fig. 3). However, the correlation was markedly stronger at low pH (R^2^ = 0.95) than at neutral pH (R^2^ = 0.61). Variation in these two quantities was also larger at low pH compared to neutral pH (Fig. 3).

At a low pH of the assay medium, *r*_0_ and *α* showed lower variation for the LpH populations compared to the ANC and NpH populations (Fig. 4 panels A-B and I-J; see also Tab. S7 in Supporting Information section S8). We did not detect differences in terms of equilibrium population density (*K*). At a neutral pH of the assay medium, we did not detect differences in variation for the intrinsic rate of increase (*r*_0_), slightly more variation in equilibrium population density (*K*), and strongly higher variation in competitive ability (*α*) of both the LpH and NpH populations compared to the ANC. Note that despite the high relative importance of the evolution variables (Evolved (general evolutionary change), LpH and NpH) for *r*_0_ at neutral pH, the effect size associated with these variables was close to zero. The high relative importance stems from the differences in how the different genotypes responded to the pH treatments, which was captured in the random effects (Fig. 4 panels C-D and K-L). In summary, we found a correlation between *r*_0_ and *α* both at low and neutral pH and found that LpH populations converged in life-history strategy, in the sense that LpH populations became more similar in life-history strategy compared to the ANC populations.

### Density-dependent fitness

While the evolutionary shifts of the individual population growth parameters were highly variable as described above, we found that under low pH of the assay medium these different changes led to an increase in the overall density-dependent fitness of the LpH populations compared to the ANC population (see also Tab. S8 in the Supporting Information section S8). No such increase in density-dependent fitness was observed for the NpH population compared to the ANC populations (see also Supporting information section S9). In both the ANC and LpH populations, density-dependent fitness increased with the intrinsic rate of increase (*r*_0_). The smaller range of *r*_0_- and *α*-values for the LpH population (Fig. 5 C and Fig. 4 panels A,B and I,J) shows the convergence of *r*_0_ discussed above. As exemplified in Fig. 5A-B, density-dependent fitness can increase whether *r*_0_ increases or decreases due to correlated changes in competitive ability *α*. In ancestral populations where the intrinsic rate of increase (*r*_0_) was initially high (Fig. 5A), competitive ability (*α*) was also high due to the strong correlation between *α* and *r*_0_. Consequently density regulation acted strongly in these populations, leading to very slow population growth (*r*) under high density conditions. Given that densities were typically high during the evolution experiment (Fig. 1; Fig. 5A), lowering *r*_0_ allowed for increased growth at higher densities and hence an increase in density-dependent fitness. If *r*_0_ was initially very low (Fig. 5B), density regulation did not act very strongly, because competitive ability (*α*) was also very low, and as a population’s intrinsic rate of increase (*r*_0_) became higher, the population’s fitness increased for all density values, leading to an increase in density-dependent fitness as well. In essence, we found that the observed convergence in life-history traits led to an average increase in density-dependent fitness at low pH for the LpH populations.

## Discussion

In this experiment, we investigated the evolutionary response of the model protist *Tetrahymena thermophila* to pH stress under high population densities. Instead of maximizing the intrinsic rate of increase (*r*_0_) we found that evolution of four different genotypes under low pH and high population density led to a convergence of life-history strategy, that is, genotypes became more similar in life-history strategy (see below). This observation stems, on the one hand, from the high population density (demography) the populations experienced during our experiment, and, on the other hand, from the genetic architecture of life-history traits, where we found that intrinsic rate of increase (*r*_0_) and competitive ability (*α*) were positively correlated, especially under stressful conditions.

Evolution can help populations adapt to changing environments (Kawecki and Ebert, 2004). Depending on the rate and severity of such change, populations need to respond quickly, as they may otherwise be driven to extinction. Past experiments have demonstrated that evolution can lead to adaptation to an abiotic stressor within few generations (Bell and Gonzalez, 2011; Padfield et al., 2016; Harmand et al., 2018). However, evidence from experimental evolution of such adaptation to pH stress remains relatively limited in many species, and is still more commonly studied using comparative work (Reusch and Boyd, 2013; Stillman and Paganini, 2015, with the notable exception of bacterial evolution experiments, as discussed above). Our results show that populations of the freshwater protist *T. thermophila* can adapt to such stress, even under conditions of strong competition due to high population densities.

Whereas our finding that evolution can alter population performance under abiotic stress agrees with the existing literature (Leimu and Fischer, 2008; Fraser et al., 2011; Kelly and Hofmann, 2013), our results on the direction of evolution were less expected. Specifically, the observed evolutionary changes in the intrinsic rates of increase (*r*_0_, Fig. 2) showed opposite directions depending on the genetic background. Many evolution experiments are conducted by serially transferring populations into fresh medium (for examples see Lenski and Travisano, 1994; Bell and Gonzalez, 2011; Bono et al., 2017). In such experiments, population densities are low during much of the period of evolution, or at least a distinct phase of selection happens under low density conditions. Under these demographic condition, selection mainly acts on the intrinsic rate of increase (*r*_0_) to maximize fitness (Mueller and Ayala, 1981). In contrast, although we use a similar approach of propagating our populations in this experiment, population densities were kept much higher (always above 50 % of population equilibrium density), leading to strongly different demographic conditions. A growing body of work on eco-evolutionary dynamics and feedbacks (Pelletier et al., 2009; Hendry, 2016) shows that it is important to consider the ecological context, here, the demographic conditions, under which evolution occurs.

This ecological context may affect how selection acts and thus alter evolutionary trajectories. Our results show that when populations evolve under high population densities, we do not find generally increased intrinsic rates of increase (*r*_0_). We suggest that this pattern is driven by the combination of genetic architecture, that is, the linkage between intrinsic rate of increase (*r*_0_) and competitive ability (*α*), constraining evolutionary trajectories (Fig. 3), and by selection for maximizing fitness under pH stress (abiotic conditions) and high population density (biotic factor). Firstly, evolution is constrained in the sense that the intrinsic rate of increase (*r*_0_) is positively correlated with competitive ability (*α*; see also Mueller and Ayala, 1981; Reznick et al., 2002; Fronhofer et al., 2018, for a different view see Joshi et al. 2001). This implies that fast growing genotypes will compete more strongly within the population than slow growing genotypes for available resources when densities increase, which is expected to slow down population growth rate at higher densities.

This slowdown in population growth rate (*r*) can clearly be seen in Fig. 5 (and Fig. S4 in the Supporting Information section S6), where genotypes that show initially a high intrinsic rate of increase (*r*_0_; high intercept) also show a strong density-dependent decrease in population growth rate (strong curvature). In contrast, populations with lower *r*_0_ show less steep declines in population growth rate. Secondly, since stress associated with low pH strongly decreased population growth rates, LpH populations experienced more difficulty to recover in population size after each medium replacement event compared to NpH populations, and hence were subject to stronger selection for increased population growth. Given that the demographic conditions were such that populations had to grow starting from 50 % of the equilibrium population density, we expect selection to lead to a maximization of population growth rate (*r*) under these specific densities experienced during evolution, that is, a maximization of density-dependent fitness (as shown in Fig. 5C).

Of course, populations may sometimes undergo quasi density-independent growth, for example during range shifts or repeated colonization and extinction events. However, whenever densities are high, growth will be density-dependent. This will often be the case in established populations, which are expected to fluctuate around their equilibrium population density. For example, environmental shifts (acid rain or temperature shifts, for instance) could lead to local changes affecting already well-established populations. As shown in our experiment, adaptation to abiotic stress under such demographic conditions can strongly affect trajectories of evolution, leading to complex evolutionary changes when populations simultaneously need to adapt to abiotic and biotic stress. In addition, as in our experiment, the direction of the evolutionary trajectory may depend on the starting conditions, and populations with different genetic backgrounds may evolve differently. We speculate here that under these high population density conditions, we can observe convergent evolution in life-history strategy, whereas under low population density conditions, we may instead expect parallel evolution where all populations shift their intrinsic rate of increase (*r*_0_) upwards at low pH. The term convergent evolution has however been defined multiple times (as discussed in Blount et al., 2018; Wood et al., 2005; Bolnick et al., 2018). We here follow the geometric argumentation in Bolnick et al. (2018). We thus define and will use the following terminology to describe evolutionary responses as follows: 1) Convergent evolution occurs when different populations develop more similar phenotypes during evolution, 2) divergent evolution implies that different populations develop more distinct phenotypes during evolution) and 3) parallel evolution occurs when different populations undergo phenotypic changes in the same direction during evolution. We should however also note that our results suggest that within genotypes, evolution happened in parallel, as all replicate populations underwent directional evolution towards either increased or decreased intrinsic rate of increase (*r*_0_), although over all genotypes, we observed convergence to a strategy that optimized the density-dependent fitness of populations.

In agreement with our observation that evolution in response to low pH may be variable, recent work has found no clear consensus on the effect of acidification on species growth rates (Kelly and Hofmann, 2013; Gattuso and Hansson, 2011, chapter 6-7). Also, shorter-term ecological experiments, despite showing a clear positive effect on photosynthesis, found that different species showed strongly differing changes in growth rates to acidification (Gattuso and Hansson, 2011, chapter 6). Similarly, longer-term evolution experiments have demonstrated that intrinsic rate of increase can either increase (Lohbeck et al., 2012; Schlüter et al., 2014) or not (Collins and Bell, 2004) for populations evolved under conditions of increased CO_2_. On a speculative note, our experiment suggests that demographic conditions may be a potential explanatory factor for such divergent results. Taking into account the demographic context and other potentially confounding eco-evolutionary interactions may help to clarify these factors in future work.

In conclusion, we found that demography affected adaptation to low pH in the protist *T. thermophila*, leading to a convergence in life-history strategies and increased high-density fitness. Our work shows that taking into account demography may be key to understanding evolutionary trajectories. In an eco-evolutionary context, quantifying density-regulation functions, that is, population growth rates as a function of population density, may be a useful way forward. Furthermore, although we observe convergent evolution in life-history strategy on a phenotypic level, it remains unclear whether this evolution is also convergent on a genetic level. As noted by Wood et al. (2005), when the genetic basis of traits is simple, convergent evolution often also has a genetic basis, but when the genetic basis is more complex, there are typically multiple paths available leading to similar phenotypic changes. An interesting avenue for future research could be to further study how the observed trade-off between intrinsic rate of increase (*r*_0_) and intraspecific competitive ability (*α*) translate to the genetic level, as we see a clear trade-off between these traits, that seems phenotypically rather constrained. If such a trade-off also exists on a genetic level, understanding this link may yield new expectations concerning convergent and parallel evolution of populations, both in presence and absence of abiotic and biotic stress.

## Acknowledgements

The authors thank Samuel Hürlemann for help with laboratory work. Funding is from the URPP Evolution in Action and the Swiss National Science Foundation, Grant No PP00P3 179089. This is publication ISEM-YYYY-XXX of the Institut des Sciences de l’Evolution – Montpellier. We would also like to acknowledge support by Swiss National Science Foundation grant 31003A 172887 and by European Research Council Advanced Grant No. 739874. We would like to thank the editorial staff of Evolution and the two referees for the helpful and constructive comments on a previous version of this manuscript.

## Author contributions

FM, FA and EAF designed the experiment. FM, AA and SM performed the experimental work. Statistical analyses were done by FM and EAF. FM, FA, AW and EAF interpreted the results. FM, FA and EAF wrote the first version of the manuscript and all authors commented the final version.

