## Supporting Information for "Evolution under pH stress and high population densities leads to increased density-dependent fitness in the protist *Tetrahymena thermophila*"

#### S1 Relationship between HCl and pH

| Amount HCl added (μL) | Total amount HCl (μL) | pH |
|---|---|---|
| 0 | 0 | 6.61 |
| 40 | 40 | 6.56 |
| 160 | 200 | 6.31 |
| 200 | 400 | 5.97 |
| 200 | 600 | 5.60 |
| 200 | 800 | 5.27 |
| 200 | 1000 | 5.0 |
| 200 | 1200 | 4.82 |
| 200 | 1400 | 4.64 |
| 200 | 1600 | 4.50 |
| 200 | 1800 | 4.37 |
| 200 | 2000 | 4.24 |
| 50 | 2050 | 4.21 |
| 50 | 2100 | 4.19 |
| 50 | 2150 | 4.15 |
| 50 | 2200 | 4.12 |
| 50 | 2250 | 4.11 |
| 50 | 2300 | 4.08 |
| 50 | 2350 | 4.06 |
| 50 | 2400 | 4.03 |
| 50 | 2450 | 4.00 |
| 100 | 2550 | 3.95 |
| 100 | 2650 | 3.89 |

Table S1: Table showing the amount of 1M HCl added to 100mL of the SSP medium, and the corresponding measured pH.

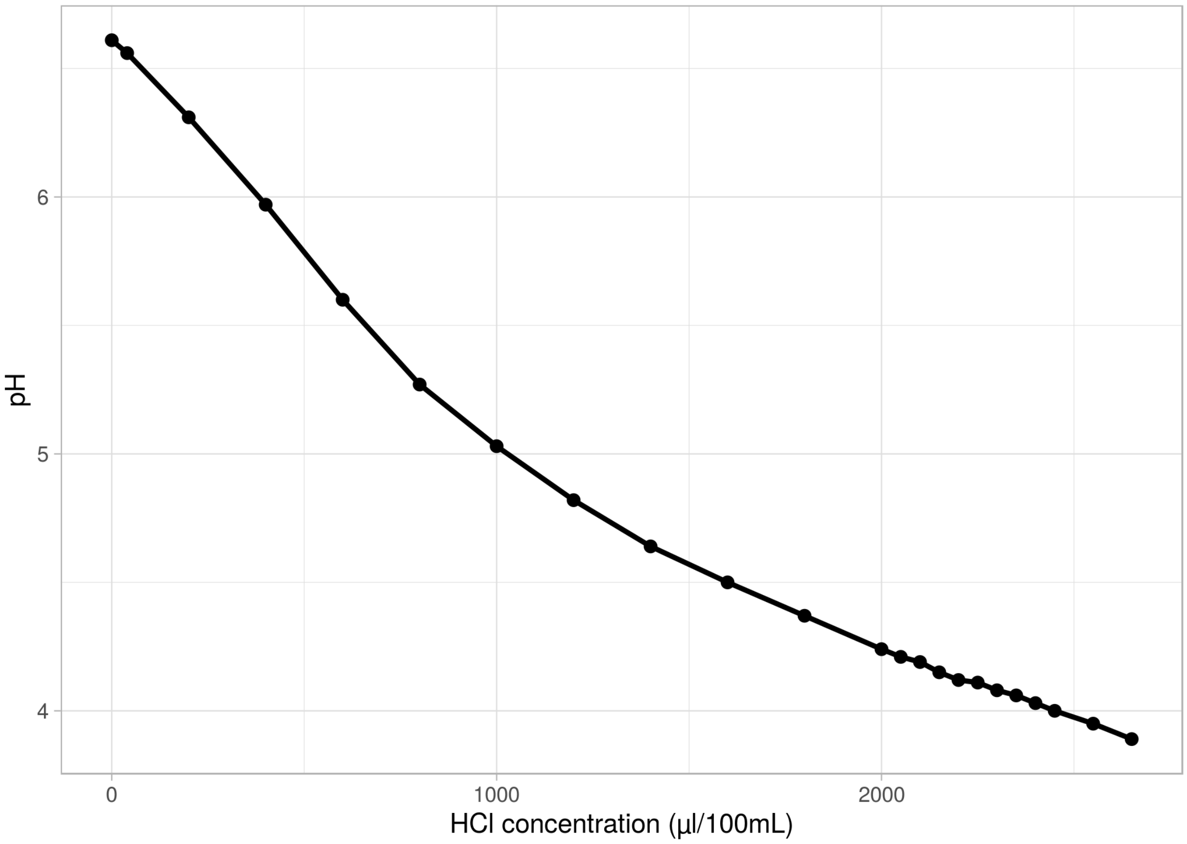


Figure S1: Measured pH (y-axis) as a function of HCl concentration (x-axis).

#### S2 PCA plot showing genotype differences

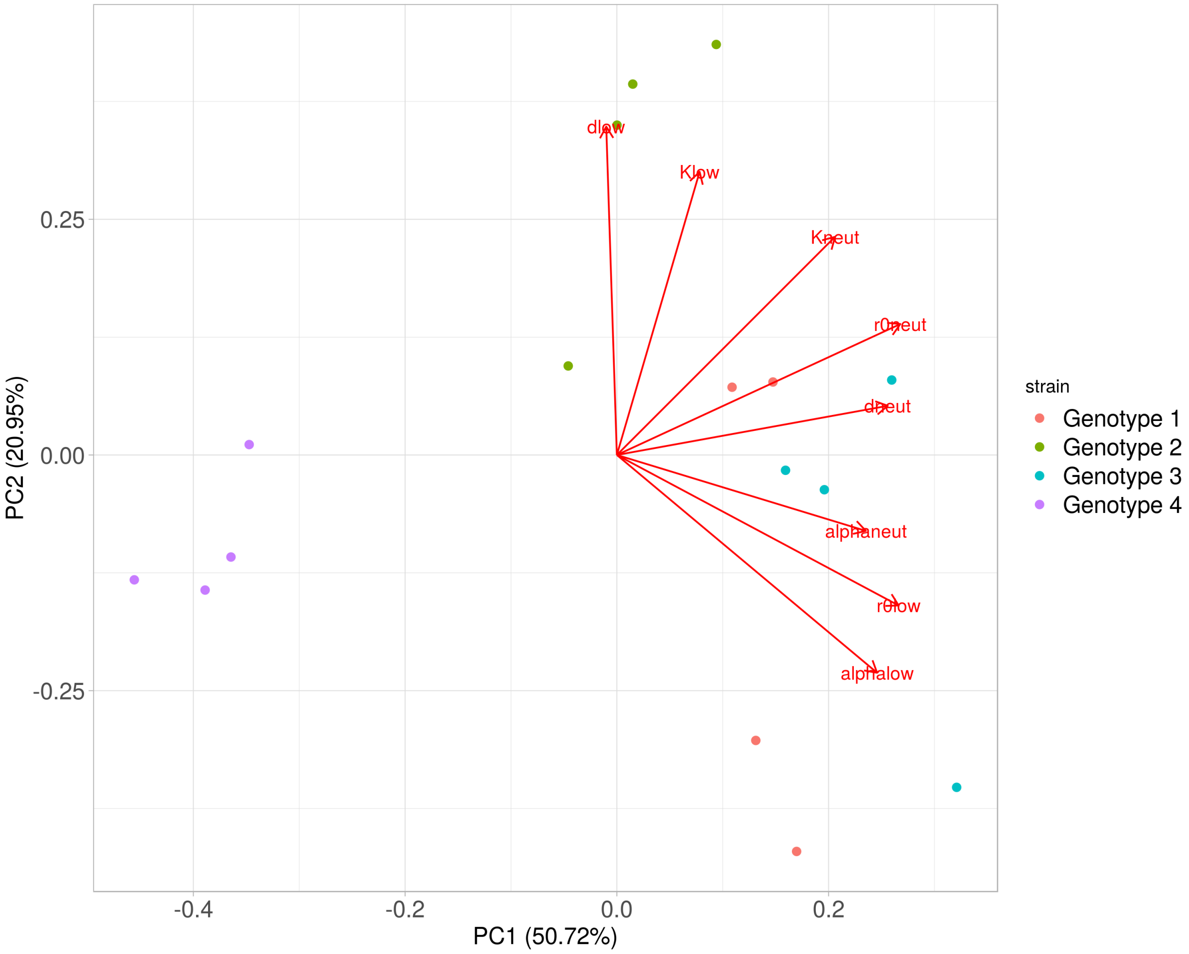


Figure S2: PCA of the ancestral genotypes, using the population growth parameters (intrinsic rate of increase ($r_0$), competitive ability ($\alpha$), mortality ($d$) and equilibrium population density ($K$)) measured both at low (4.5) and neutral (6.5) pH. Genotypes are coloured in red (genotype 1 – B2086.2), green (genotype 2 – CU427.4), blue (genotype 3 – CU428.2) and purple (genotype 4 – SB3539).

#### S3 Video analysis script

```
###########################################################################
# R script for analysing video files with BEMOVI (www.bemovi.info)
rm(list=ls())
# load package
library(devtools)
install_github("efronhofer/bemovi", ref="experimental")
library(bemovi)

###########################################################################
# VIDEO PARAMETERS

# video frame rate (in frames per second)
fps <- 25
# length of video (in frames)
total_frames <- 500

# measured volume (in microliter)
measured_volume <- 34.4 # for Leica M205 C with 1.6 fold magnification,
Sample  height 0.5 mm and Hamamatsu Orca Flash 4

# size of a pixel (in micrometer)
pixel_to_scale <- 4.05 # for Leica M205 C with 1.6 fold magnification,
Sample  height 0.5 mm and Hamamatsu Orca Flash 4

# specify video file format (one of "avi","cxd","mov","tiff")
# bemovi only works with avi and cxd. other formats are reformated to avi below
video.format <- "cxd"

```

```
# setup
difference.lag <- 10
thresholds <- c(10,255) # don't change the second value
#thresholds <- c(50,255)

########################################################################
# FILTERING PARAMETERS
# min and max size: area in pixels
particle_min_size <- 5
particle_max_size <- 1000

# number of adjacent frames to be considered for linking particles
trajectory_link_range <- 3
# maximum distance a particle can move between two frames
trajectory_displacement <- 16

# these values are in the units defined by the parameters above: fps (seconds),
#measured_volume (microliters) and pixel_to_scale (micometers)
filter_min_net_disp <- 25
filter_min_duration <- 1
filter_detection_freq <- 0.1
filter_median_step_length <- 3

########################################################################
# MORE PARAMETERS (USUALLY NOT CHANGED)

# set paths to ImageJ and particle linker standalone
IJ.path <- "/home/felix/bin/ImageJ"
to.particlelinker <- "/home/felix/bin/ParticleLinker"
```

```
# directories and file names
to.data <- paste(getwd(),"/",sep="")
video.description.folder <- "0_video_description/"
video.description.file <- "video_description.txt"
raw.video.folder <- "1_raw/"
particle.data.folder <- "2_particle_data/"
trajectory.data.folder <- "3_trajectory_data/"
temp.overlay.folder <- "4a_temp_overlays/"
overlay.folder <- "4_overlays/"
merged.data.folder <- "5_merged_data/"
ijmacs.folder <- "ijmacs/"

# RAM allocation
memory.alloc <- c(60000)

# RAM per particle linker instance
memory.alloc.perLinker <- c(10000)

######################################################################
# VIDEO ANALYSIS

# identify particles
locate_and_measure_particles(to.data, raw.video.folder, particle.data.folder,
difference.lag, thresholds, min_size = particle_min_size, max_size =
particle_max_size, IJ.path, memory.alloc)

# link the particles
link_particles(to.data, particle.data.folder, trajectory.data.folder, linkrange =
```

```
trajectory_link_range, disp = trajectory_displacement, start_vid = 1, memory =
memory.alloc, memory_per_linkerProcess = memory.alloc.perLinker)

# merge info from description file and data
merge_data(to.data, particle.data.folder, trajectory.data.folder,
video.description.folder, video.description.file, merged.data.folder)

# load the merged data
load(paste0(to.data, merged.data.folder, "Master.RData"))

# filter data: minimum net displacement, their duration, the detection
#frequency and the median step length
trajectory.data.filtered <- filter_data(trajectory.data, filter_min_net_disp,
filter_min_duration, filter_detection_freq, filter_median_step_length)

# summarize trajectory data to individual-based data
morph_mvt <- summarize_trajectories(trajectory.data.filtered, calculate.median=F,
write = T, to.data, merged.data.folder)

# get Sample  level info
summarize_populations(trajectory.data.filtered, morph_mvt, write=T, to.data,
merged.data.folder, video.description.folder, video.description.file, total_frames)

# create overlays for validation
create_overlays(trajectory.data.filtered, to.data, merged.data.folder,
raw.video.folder, temp.overlay.folder, overlay.folder, 2048, 2048,
difference.lag, type = "label", predict_spec = F, IJ.path,
contrast.enhancement = 1, memory = memory.alloc)
```

#### S4 Model priors for fitting Bayesian evolution models

Models were always fit with intercept estimates that corresponded to the genotype mean, but with a broad enough standard deviation so the model was not constrained too much. We ran all models with a long chain length (warmup = 40,000 iterations, chain = 160,000 iterations).

##### S4.1 Models for $r_0$

Models were fit with the following priors:

- Intercepts were fit following a normal distribution with a genotype specific mean and standard deviation 0.5: int $\sim$ normal($\mu_{\text{genotype}}$, 0.5)
- All other fixed effects were modelled using a normal distribution with mean 0 and sdtandard deviation 1: effect $\sim$ normal(0, 1)
- Model standard deviation was fit using a cauchy distribution: sigma $\sim$ cauchy(0, 1)

##### S4.2 Models for $K$

Models were fit with the following priors:

- Intercepts were fit following a normal distribution with a genotype specific mean and standard deviation 0.5: int $\sim$ normal($\mu_{\text{genotype}}$, 0.5)
- All other fixed effects were modelled using a normal distribution with mean 0 and sdtandard deviation 1: effect $\sim$ normal(0, 1)
- Model standard deviation was fit using a cauchy distribution: sigma $\sim$ cauchy(0, 1)

##### S4.3 Models for $\alpha$

Models were fit with the following priors:

- Intercepts were fit following a normal distribution with a genotype specific mean and standard deviation 0.5: int $\sim$ normal($\mu_{\text{genotype}}$, 0.5)

- All other fixed effects were modelled using a normal distribution with mean 0 and sdtandard deviation 1: effect $\sim$ normal(0, 1)

- Model standard deviation was fit using a cauchy distribution: sigma $\sim$ cauchy(0, 1)

### S5 Correlation $r_0$-$\alpha$

#### S5.1 Model code

```
#Stan code
{
model <- "
 // Pearson Correlation
 data {
 int<lower=1> n;
 vector[2] x_obs[n];
 vector[2] x_sd[n];
 }
 parameters {
 vector[2] mu;
 vector<lower=0>[2] lambda;
 real<lower=-1,upper=1> r;
 vector[2] x_est[n];

 }
 transformed parameters {
 vector<lower=0>[2] sigma;
 cov_matrix[2] T;

 // Reparameterization
 sigma[1] = sqrt(lambda[1]);
 sigma[2] = sqrt(lambda[2]);
 T[1,1] = square(sigma[1]);
 T[1,2] = r * sigma[1] * sigma[2];
 T[2,1] = r * sigma[1] * sigma[2];
 T[2,2] = square(sigma[2]);
```

```
}
model {
// Priors
mu ~ normal(0, 10);
lambda ~ normal(0,1);
r ~ normal(0,1);

// Data
x_est ~ multi_normal(mu, T);

for (i in 1:n){
x_obs[i][1] ~ normal(x_est[i][1], x_sd[i][1]);
x_obs[i][2] ~ normal(x_est[i][2], x_sd[i][2]);
}

}"
}
```

##### S5.2 Correlation test for individual groups

| Origin | $R^2$ (low pH of assay medium) | $R^2$ (low pH of assay medium) |
|---|---|---|
| **ANC** | 0.97 | 0.33 |
| **LpH** | 0.49 | 0.42 |
| **NpH** | 0.97 | 0.45 |

Table S2: Correlation (median $R^2$) between intrinsic rate of increase $r_0$ and competitive ability $\alpha$ for the ANC, LpH and NpH populations, measured either at low pH of the assay medium or neutral pH of the assay medium. Numbers rounded to 2 decimal digits.

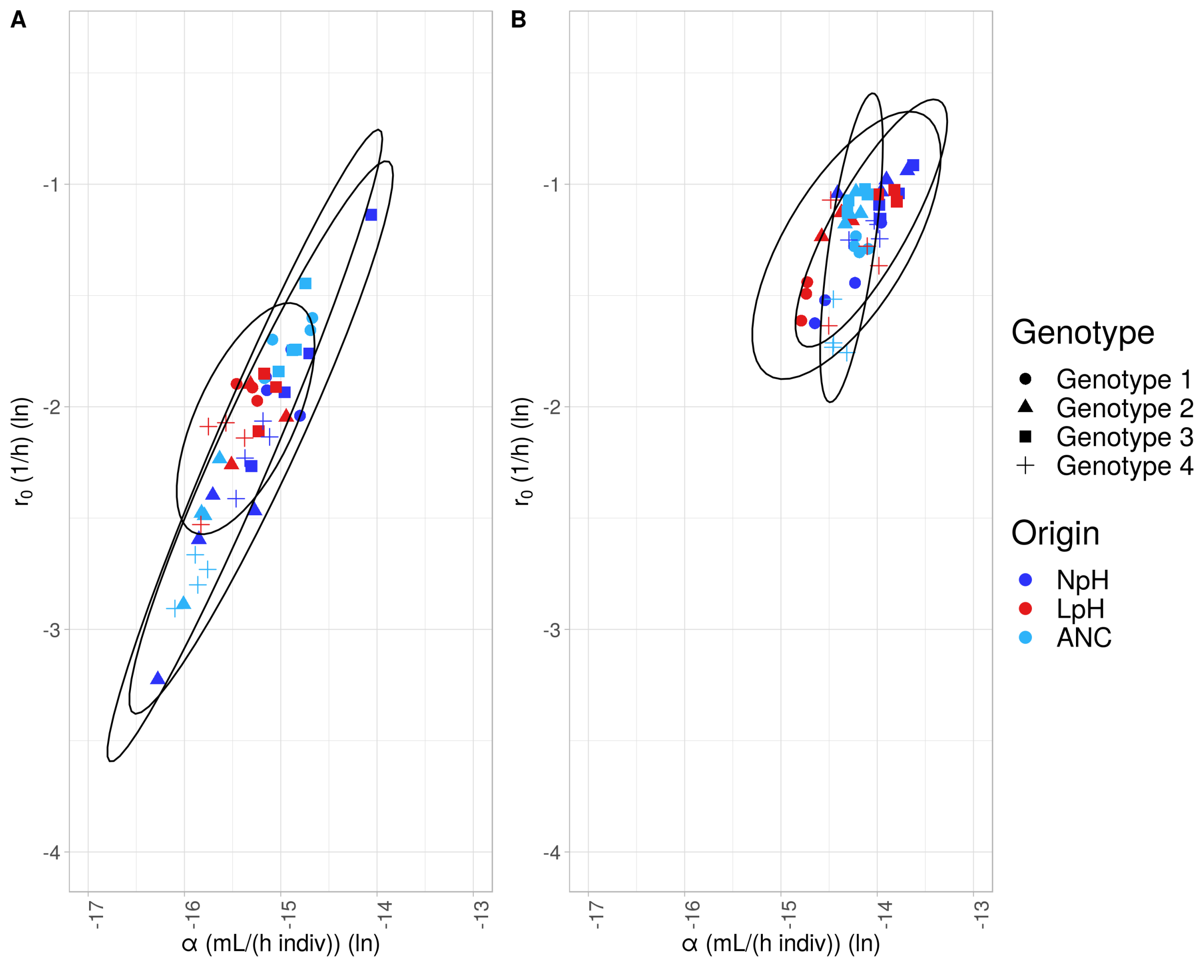


Figure S3: Correlation between the intrinsic rate of increase ($r_0$) and competitive ability ($\alpha$) at low pH (A), and at neutral pH (B) of the assay medium, separate for the ANC, LpH and NpH populations. Symbols represent the different genotypes (see legend); Light blue = ANC (ancestor populations), dark blue = NpH (populations evolved under neutral pH conditions), red = LpH (populations evolved under low pH conditions). Ellipses represents 95% probability intervals.

868 ## S6 Density regulation functions

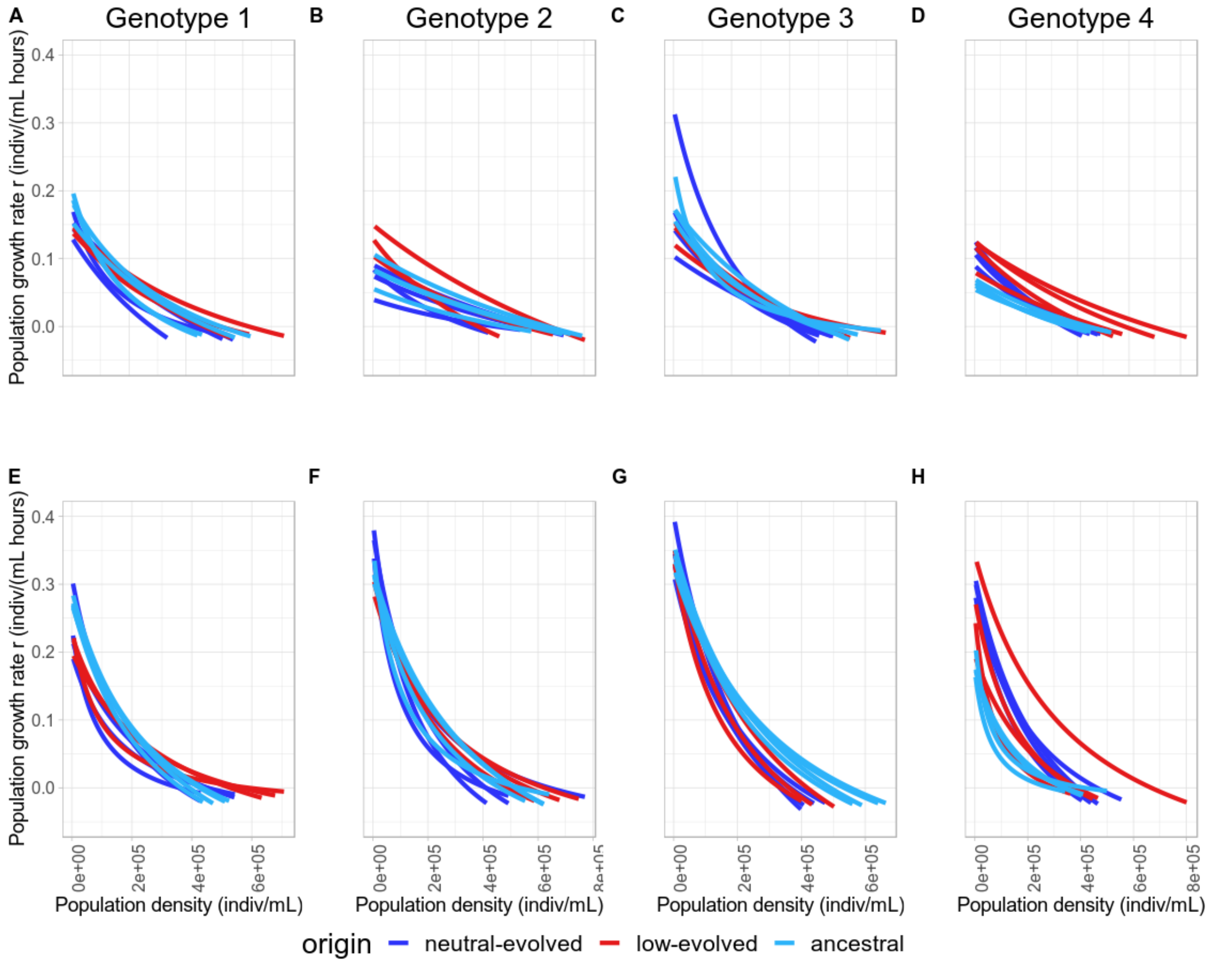


Figure S4: Density-regulation functions of the four genotypes. The y-axis depicts the population growth rate ($r$), x-axis shows the density of the population. Red lines = LpH populations, dark blue lines = NpH populations (populations evolved under low pH conditions), light blue = ANC populations (ancestral populations). Top row is the density-regulation for low pH of the assay medium Bottom row for neutral pH of the assay medium

#### S7 Model construction tables

| | pHmedium | Evolved | LpH | NpH | Evolved*pHmedium | LpH*pHmedium | NpH*pHmedium |
|---|---|---|---|---|---|---|---|
| **Model 1** | | | | | | | |
| **Model 2** | X | | | | | | |
| **Model 3** | | X | | | | | |
| **Model 4** | | | X | | | | |
| **Model 5** | | | | X | | | |
| **Model 6** | | | X | X | | | |
| **Model 7** | X | X | | | | | |
| **Model 8** | X | X | | | | X | |
| **Model 9** | X | | X | | | | |
| **Model 10** | X | | X | | | X | |
| **Model 11** | X | | | X | | | |
| **Model 12** | X | | | X | | | X |
| **Model 13** | X | | X | X | | | |
| **Model 14** | X | | X | X | | X | |
| **Model 15** | X | | X | X | | | X |
| **Model 16** | X | | X | X | | X | X |

Table S3: The different statistical models included for the model averaging of $r_0$, $K$ and $\alpha$. Rows are the different models and columns the different factors. pHmedium = plastic effect of assay medium pH, evolved = general evolutionary shifts. LpH = evolution effect in population evolving at low pH, NpH = evolution effect in populations evolving at neutral pH. interaction effects show pH specific evolution effects for the evolution factors.

| | Evolved | LpH | NpH | Strain effects |
|---|---|---|---|---|
| **Model 1** | | | | |
| **Model 2** | X | | | |
| **Model 3** | | X | | |
| **Model 4** | | | X | |
| **Model 5** | | X | X | |
| **Model 6** | | | | X |
| **Model 7** | X | | | X |
| **Model 8** | | X | | X |
| **Model 9** | | | X | X |
| **Model 10** | | X | X | X |

Table S4: Parameters included in the different models for analyzing group differences in variation in life-history traits. Evolved = general difference between ANC and evolved populations. LpH = specific differences for populations evolved at low pH. NpH = specific differences for populations evolved at neutral pH. Genotypes effects = inclusion of random genotype intercepts.

| | Centered $r_0$ | Origin | Origin* Centered $r_0$ |
|---|---|---|---|
| **Model 1** | | | |
| **Model 2** | X | | |
| **Model 3** | | X | |
| **Model 4** | X | X | |
| **Model 5** | X | X | X |

Table S5: Parameters included in the different models for analyzing density-dependent fitness estimates. Included explanatory variables are a) Centered $r_0$: numerical variable associated with effect of the centered intrinsic rate of increase $r_0$), b) Origin: Categorical variable, with factors LpH (1) or Ancestral (0), c) Origin*Centered $r_0$: Numerical variable, interaction term between Centered $r_0$ and Origin

### S8 Relative importance tables

| | | Genotype 1 | | Genotype 2 | | Genotype 3 | | Genotype 4 | |
|---|---|---|---|---|---|---|---|---|---|
| | | **RI** | **Effect** | **RI** | **Effect** | **RI** | **Effect** | **RI** | **Effect** |
| $r_0$ | **pHmedium** | 1 | + | 1 | + | 1 | + | 1 | + |
| | **evolved** | 0.51 | - | 0.02 | + | 0.34 | - | 0.56 | + |
| | **LpH** | 0.37 | - | 0.92 | + | 0.38 | - | 0.43 | + |
| | **NpH** | 0.42 | - | 0.4 | - | 0.35 | - | 0.43 | + |
| | **evolved*pHmedium** | 0.06 | + | 0.01 | - | 0.16 | + | 0.22 | - |
| | **NpH*pHmedium** | 0.14 | + | 0.16 | + | 0.15 | + | 0.13 | +/- |
| | **LpH*pHmedium** | 0.1 | - | 0.74 | - | 0.14 | + | 0.19 | - |
| $K$ | **pHmedium** | 0.9 | - | 0.56 | - | 0.94 | - | 0.96 | - |
| | **evolved** | 0.01 | + | 0.12 | - | 0.45 | - | 0.11 | + |
| | **LpH** | 0.89 | + | 0.35 | + | 0.4 | - | 0.75 | + |
| | **NpH** | 0.49 | - | 0.42 | - | 0.55 | - | 0.38 | +/- |
| | **evolved*pHmedium** | 0 | / | 0.02 | + | 0.21 | - | 0.03 | + |
| | **NpH*pHmedium** | 0.17 | + | 0.07 | - | 0.53 | - | 0.15 | + |
| | **LpH*pHmedium** | 0.12 | + | 0.06 | + | 0.21 | - | 0.26 | - |
| $\alpha$ | **pHmedium** | 1 | + | 1 | + | 1 | + | 1 | + |
| | **evolved** | 0.03 | - | 0.16 | + | 0.28 | - | 0.16 | + |
| | **LpH** | 0.93 | - | 0.52 | + | 0.35 | - | 0.34 | +/- |
| | **NpH** | 0.34 | +/- | 0.39 | +/- | 0.31 | + | 0.69 | + |
| | **evolved*pHmedium** | 0.01 | - | 0.05 | + | 0.23 | + | 0.05 | - |
| | **NpH*pHmedium** | 0.11 | - | 0.17 | + | 0.13 | + | 0.26 | - |
| | **LpH*pHmedium** | 0.27 | - | 0.36 | - | 0.18 | + | 0.1 | + |

Table S6: Relative importance (RI) of explanatory variables for $r_0$, $K$ and $\alpha$ for the 4 different genotypes. pHmedium = plastic effect of assay medium pH, evolved = general evolutionary shifts (difference between ANC and evolved populations). LpH = evolution effect in population evolving at low pH, NpH = evolution effect in populations evolving at neutral pH. Interaction effects show pH specific evolution effects for the evolutionary origin (LpH or NpH). RI gives the relative importance of the factors, effect shows the direction of the effect (+ = positive, - = negative, +/- = different depending on model).

| | | $r_0$ | | $K$ | | $\alpha$ | |
|---|---|---|---|---|---|---|---|
| | | **RI** | **Effect** | **RI** | **Effect** | **RI** | **Effect** |
| **Low assay** | **Evolved** | 0.05 | - | 0.26 | 0 | 0.18 | - |
| **medium pH** | **LpH** | 0.92 | - | 0.29 | 0 | 0.77 | - |
| | **NpH** | 0.25 | - | 0.25 | 0 | 0.18 | - |
| **Neutral assay** | **Evolved** | 0.71 | 0 | 0.04 | 0 | 0.15 | + |
| **medium pH** | **LpH** | 0.29 | 0 | 0.7 | + | 0.86 | + |
| | **NpH** | 0.29 | 0 | 0.46 | - | 0.85 | + |

Table S7: Relative importance (RI) of evolutionary origin on trait convergence. RI gives relative importance associated with evolutionary history (evolved = general evolutionary response, LpH = evolution effect in populations evolving at low pH, NpH = evolution effect in populations evolving at neutral pH). The "Effect" column shows the direction (sign) of change.

| | **RI** | **Effect** |
|---|---|---|
| **Centered** $r_0$ | 1 | + |
| **Origin** | 1 | + |
| **Origin*Centered** $r_0$ | 0.6 | + |

Table S8: Relative importance (RI) for density-dependent fitness models. RI gives relative importance associated with a) Centered $r_0$: numerical variable associated with effect of the centered intrinsic rate of increase $r_0$), b) Origin: Categorical variable, with factors LpH (1) or Ancestral (0), c) Origin*Centered $r_0$: Numerical variable, interaction term between Centered $r_0$ and Origin. The "Effect" column shows the direction (sign) of change.

#### S9 Density-dependent fitness of the NpH populations

We repeated the analysis for density-dependent fitness for the NpH populations. We calculated the population growth rate ($r$) for the NpH and for ANC populations over all observed population densities during the evolution experiment and integrated over these values to calculate a weighted density-dependent fitness estimate. We then used Bayesian models to fit these density-dependent fitness values as a function of a) population origin (ANC = 0 or NpH = 1), b) centered intrinsic rate of increase ($r_0$), and c) an interaction term between $r_0$ and population origin. Centered $r_0$ represents the intrinsic rate of increase, rescaled to have its mean at zero, and was calculated by subtracting the mean $r_0$ from all $r_0$ values. In this analysis, we also included a random intercept for the different genotypes. We fit all five models, starting from the intercept model to the full interaction model. Subsequently, we ranked these models using the WAIC criterion.

Only the full interaction model was considered. Below, in Tab. S9 are listed the summary statistics of the model (mean estimates, standard deviation and 95 %-confidence intervals). Note how the density-dependent fitness of NpH populations does on average not differ (0 is included in the 95 %-confidence interval; Tab. S9) and even has the tendency to be slightly lower than the ANC populations. Note also how the relation between density-dependent fitness and centered $r_0$ changes from positive in ANC to negative in NpH (Tab. S9; Fig. S5).

| **Variable** | **Mean estimate** | **Standard deviation** | **95 %-confidence interval** |
|---|---|---|---|
| **Intercept** | 2.43 | 0.92 | (0.56, 4.18) |
| **Genotype 1 (random intercept)** | -1.34 | 0.97 | (-3.25, 0.56) |
| **Genotype 2 (random intercept)** | -0.08 | 0.97 | (-1.98, 1.84) |
| **Genotype 3 (random intercept)** | -1.10 | 0.97 | (-3.02, 0.79) |
| **Genotype 4 (random intercept)** | 2.63 | 0.99 | (0.70, 4.55) |
| **Centered** $r_0$ | 19.90 | 4.98 | (10.08, 29.50) |
| **Origin** | -0.72 | 0.40 | (-1.50, 0.07) |
| **centered** $r_0$ *** origin** | -31.80 | 5.45 | (-42.22, -20.86) |

Table S9: Mean estimate, standard deviation and 95 %-confidence intervals for the fixed and random factors included in the best (and single considered model) from WAIC comparison. All numbers are rounded to 2 decimal digits.

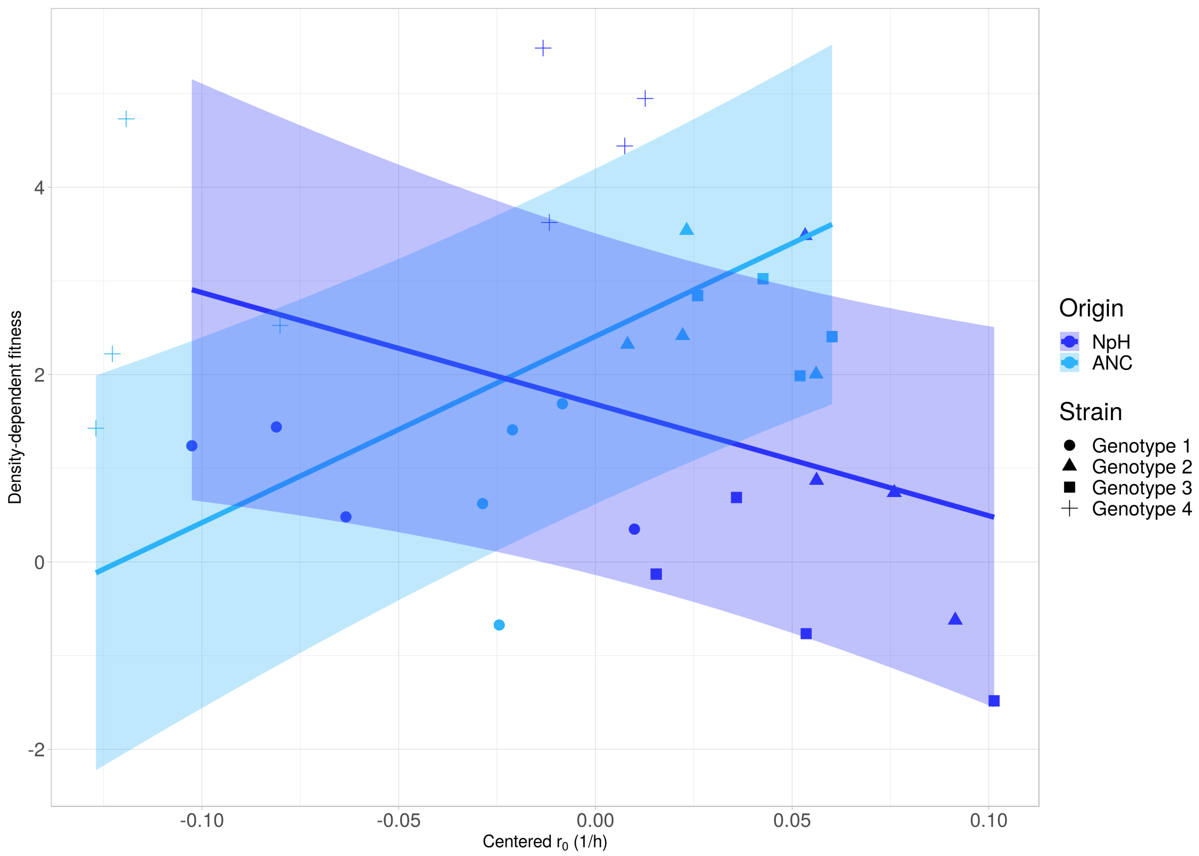


Figure S5: Density-dependent fitness depending on the (centered) intrinsic rate of increase ($r_0$). Symbols correspond to data from NpH (dark blue) or ANC (light blue) populations (shape represents genotype, see legend). Lines and shaded areas represent the weighted posterior predictions and the 95% probability intervals for the four genotypes.

#### S10 Population growth assessment: raw data and posterior predictions

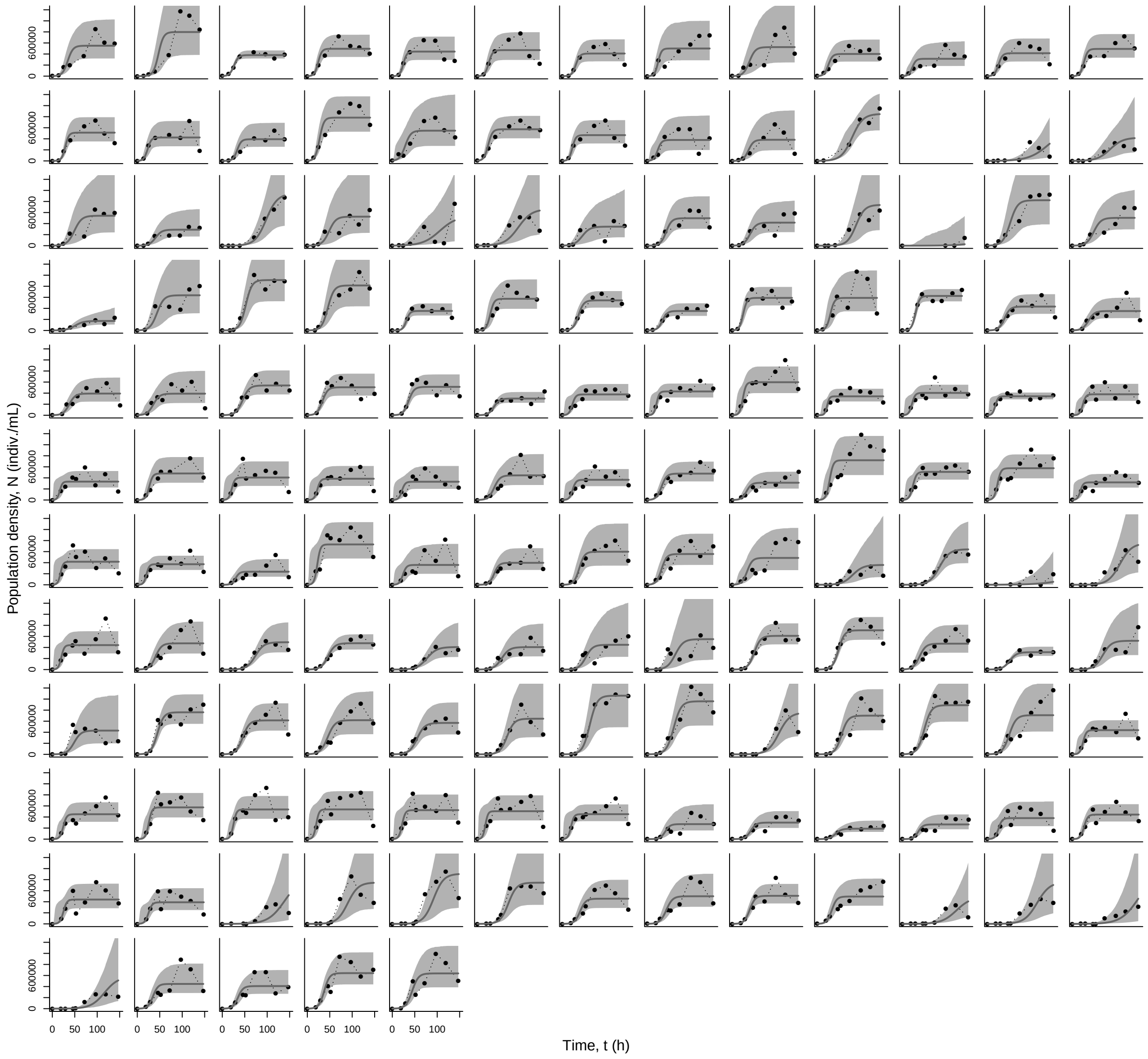


Figure S6: Individual subplots show the population density measurements and model predictions for a single population during the population growth assessments. Points are measured population densities. Lines show the mean model predictions, and shaded areas the 95 % confidence intervals of the posterior predictions.

| Replicate | $\log(r_o)$ | | $\log(K)$ | | $\log(d)$ | | $\log(\alpha)$ | |
|---|---|---|---|---|---|---|---|---|
| | mean | sd | mean | sd | mean | sd | mean | sd |
| Sample 0001 | -1.033 | 0.276 | 12.925 | 0.123 | -2.480 | 0.765 | -13.958 | 0.291 |
| Sample 0002 | -0.979 | 0.318 | 12.925 | 0.195 | -1.623 | 1.110 | -13.904 | 0.393 |
| Sample 0003 | -1.041 | 0.239 | 13.372 | 0.144 | -2.321 | 0.876 | -14.413 | 0.270 |
| Sample 0004 | -0.939 | 0.267 | 12.748 | 0.215 | -1.660 | 1.107 | -13.687 | 0.363 |

| | | | | | | | | |
|---|---|---|---|---|---|---|---|---|
| Sample 0005 | -1.092 | 0.291 | 12.888 | 0.157 | -1.617 | 1.069 | -13.980 | 0.336 |
| Sample 0006 | -0.914 | 0.301 | 12.714 | 0.120 | -1.164 | 0.979 | -13.628 | 0.338 |
| Sample 0007 | -1.155 | 0.277 | 12.811 | 0.214 | -1.537 | 1.097 | -13.966 | 0.367 |
| Sample 0008 | -1.041 | 0.346 | 12.735 | 0.200 | -1.424 | 1.069 | -13.776 | 0.423 |
| Sample 0009 | -1.251 | 0.235 | 13.043 | 0.196 | -1.911 | 1.037 | -14.293 | 0.313 |
| Sample 0010 | -1.164 | 0.340 | 12.867 | 0.243 | -1.479 | 1.120 | -14.031 | 0.445 |
| Sample 0011 | -1.181 | 0.336 | 12.810 | 0.245 | -1.565 | 1.129 | -13.991 | 0.441 |
| Sample 0012 | -1.245 | 0.393 | 12.729 | 0.276 | -1.799 | 1.159 | -13.974 | 0.523 |
| Sample 0013 | -1.624 | 0.254 | 13.024 | 0.225 | -2.576 | 1.083 | -14.648 | 0.333 |
| Sample 0014 | -1.173 | 0.329 | 12.787 | 0.217 | -1.766 | 1.104 | -13.959 | 0.421 |
| Sample 0015 | -1.521 | 0.267 | 13.021 | 0.193 | -2.096 | 1.103 | -14.542 | 0.330 |
| Sample 0016 | -1.443 | 0.279 | 12.787 | 0.226 | -2.878 | 1.052 | -14.230 | 0.327 |
| Sample 0017 | -1.234 | 0.301 | 13.342 | 0.239 | -1.999 | 1.152 | -14.577 | 0.397 |
| Sample 0018 | -1.163 | 0.289 | 13.100 | 0.152 | -1.797 | 1.008 | -14.263 | 0.332 |
| Sample 0019 | -1.127 | 0.298 | 13.244 | 0.169 | -1.898 | 1.092 | -14.371 | 0.337 |
| Sample 0020 | -1.078 | 0.363 | 12.718 | 0.201 | -1.812 | 1.118 | -13.797 | 0.436 |
| Sample 0021 | -1.046 | 0.310 | 12.945 | 0.174 | -1.286 | 1.043 | -13.991 | 0.379 |
| Sample 0022 | -1.025 | 0.317 | 12.797 | 0.192 | -1.486 | 1.074 | -13.822 | 0.394 |
| Sample 0023 | -1.366 | 0.291 | 12.617 | 0.286 | -2.782 | 1.176 | -13.983 | 0.397 |
| Sample 0024 | -1.071 | 0.298 | 13.411 | 0.192 | -1.678 | 1.005 | -14.482 | 0.376 |
| Sample 0025 | -1.278 | 0.354 | 12.825 | 0.258 | -1.967 | 1.182 | -14.104 | 0.457 |
| Sample 0026 | -1.636 | 0.196 | 12.868 | 0.237 | -1.904 | 1.183 | -14.504 | 0.321 |
| Sample 0027 | -1.613 | 0.250 | 13.176 | 0.241 | -1.968 | 1.173 | -14.789 | 0.363 |
| Sample 0028 | -1.492 | 0.249 | 13.245 | 0.183 | -2.404 | 1.037 | -14.737 | 0.295 |
| Sample 0029 | -1.440 | 0.304 | 13.287 | 0.272 | -3.282 | 1.009 | -14.727 | 0.377 |
| Sample 0030 | -2.466 | 0.210 | 12.806 | 0.383 | -2.138 | 1.392 | -15.272 | 0.473 |
| Sample 0031 | -2.397 | 0.090 | 13.309 | 0.224 | -1.304 | 1.170 | -15.706 | 0.257 |
| Sample 0032 | -3.225 | 0.453 | 13.051 | 0.502 | -2.579 | 1.665 | -16.277 | 0.742 |
| Sample 0033 | -2.595 | 0.168 | 13.255 | 0.381 | -1.835 | 1.259 | -15.850 | 0.432 |

| | | | | | | | | |
|---|---|---|---|---|---|---|---|---|
| Sample 0034 | -1.138 | 0.360 | 12.923 | 0.258 | -1.574 | 1.137 | -14.061 | 0.471 |
| Sample 0035 | -1.760 | 0.251 | 12.945 | 0.267 | -2.018 | 1.202 | -14.705 | 0.388 |
| Sample 0036 | -2.267 | 0.101 | 13.037 | 0.228 | -1.251 | 1.164 | -15.304 | 0.260 |
| Sample 0037 | -1.935 | 0.116 | 13.025 | 0.149 | -1.790 | 1.005 | -14.960 | 0.194 |
| Sample 0038 | -2.413 | 0.120 | 13.051 | 0.266 | -1.734 | 1.264 | -15.464 | 0.307 |
| Sample 0039 | -2.135 | 0.145 | 12.980 | 0.248 | -1.622 | 1.244 | -15.116 | 0.304 |
| Sample 0040 | -2.064 | 0.182 | 13.120 | 0.298 | -2.019 | 1.260 | -15.184 | 0.369 |
| Sample 0041 | -2.230 | 0.259 | 13.141 | 0.385 | -2.006 | 1.333 | -15.372 | 0.489 |
| Sample 0042 | -1.926 | 0.110 | 13.219 | 0.167 | -1.195 | 1.053 | -15.144 | 0.211 |
| Sample 0043 | -1.866 | 0.137 | 13.287 | 0.214 | -1.153 | 1.111 | -15.153 | 0.270 |
| Sample 0044 | -1.742 | 0.194 | 13.148 | 0.188 | -2.686 | 0.956 | -14.890 | 0.255 |
| Sample 0045 | -2.040 | 0.082 | 12.760 | 0.138 | -1.254 | 1.118 | -14.800 | 0.164 |
| Sample 0046 | -2.260 | 0.145 | 13.251 | 0.284 | -1.831 | 1.237 | -15.512 | 0.331 |
| Sample 0047 | -2.046 | 0.229 | 12.897 | 0.327 | -1.579 | 1.285 | -14.943 | 0.414 |
| Sample 0048 | -1.899 | 0.128 | 13.414 | 0.188 | -1.032 | 1.108 | -15.313 | 0.239 |
| Sample 0049 | -1.911 | 0.182 | 13.138 | 0.228 | -1.445 | 1.140 | -15.049 | 0.312 |
| Sample 0050 | -1.851 | 0.198 | 13.322 | 0.210 | -2.562 | 1.037 | -15.173 | 0.292 |
| Sample 0051 | -2.110 | 0.143 | 13.121 | 0.236 | -1.309 | 1.172 | -15.231 | 0.292 |
| Sample 0052 | -2.140 | 0.151 | 13.235 | 0.293 | -1.503 | 1.189 | -15.375 | 0.336 |
| Sample 0053 | -2.089 | 0.153 | 13.663 | 0.296 | -1.401 | 1.157 | -15.751 | 0.344 |
| Sample 0054 | -2.072 | 0.154 | 13.498 | 0.270 | -1.428 | 1.156 | -15.570 | 0.325 |
| Sample 0055 | -2.529 | 0.117 | 13.300 | 0.319 | -1.484 | 1.200 | -15.829 | 0.350 |
| Sample 0056 | -1.973 | 0.142 | 13.271 | 0.252 | -1.426 | 1.151 | -15.244 | 0.302 |
| Sample 0057 | -1.897 | 0.144 | 13.561 | 0.181 | -1.877 | 1.034 | -15.458 | 0.246 |
| Sample 0058 | -1.913 | 0.174 | 13.381 | 0.231 | -2.170 | 1.137 | -15.295 | 0.300 |
| Sample 0059 | -1.131 | 0.213 | 13.041 | 0.132 | -1.799 | 0.976 | -14.172 | 0.249 |
| Sample 0060 | -1.034 | 0.275 | 13.187 | 0.161 | -2.696 | 0.948 | -14.221 | 0.277 |
| Sample 0061 | -1.178 | 0.255 | 13.156 | 0.194 | -1.470 | 1.057 | -14.335 | 0.339 |
| Sample 0062 | -1.134 | 0.214 | 13.155 | 0.175 | -1.438 | 1.084 | -14.289 | 0.283 |

| | | | | | | | | |
|---|---|---|---|---|---|---|---|---|
| Sample 0063 | -1.072 | 0.300 | 13.224 | 0.184 | -1.608 | 1.076 | -14.297 | 0.366 |
| Sample 0064 | -1.122 | 0.282 | 13.188 | 0.163 | -1.569 | 1.019 | -14.310 | 0.339 |
| Sample 0065 | -1.022 | 0.341 | 13.105 | 0.191 | -1.433 | 1.039 | -14.127 | 0.415 |
| Sample 0066 | -1.045 | 0.283 | 13.054 | 0.133 | -1.531 | 1.027 | -14.099 | 0.319 |
| Sample 0067 | -1.713 | 0.217 | 12.741 | 0.325 | -2.555 | 1.282 | -14.454 | 0.404 |
| Sample 0068 | -1.733 | 0.149 | 12.728 | 0.212 | -2.289 | 1.143 | -14.460 | 0.258 |
| Sample 0069 | -1.757 | 0.156 | 12.560 | 0.274 | -3.296 | 0.947 | -14.316 | 0.279 |
| Sample 0070 | -1.517 | 0.158 | 12.939 | 0.256 | -3.497 | 0.861 | -14.456 | 0.263 |
| Sample 0071 | -1.290 | 0.309 | 12.804 | 0.244 | -1.599 | 1.134 | -14.095 | 0.424 |
| Sample 0072 | -1.278 | 0.235 | 12.962 | 0.180 | -1.654 | 1.090 | -14.240 | 0.305 |
| Sample 0073 | -1.234 | 0.293 | 12.988 | 0.216 | -1.755 | 1.166 | -14.221 | 0.377 |
| Sample 0074 | -1.306 | 0.269 | 12.880 | 0.199 | -1.436 | 1.121 | -14.185 | 0.352 |
| Sample 0075 | -2.888 | 0.234 | 13.123 | 0.460 | -2.134 | 1.394 | -16.010 | 0.543 |
| Sample 0076 | -2.489 | 0.173 | 13.303 | 0.363 | -1.628 | 1.241 | -15.792 | 0.412 |
| Sample 0077 | -2.477 | 0.186 | 13.346 | 0.378 | -1.665 | 1.251 | -15.822 | 0.430 |
| Sample 0078 | -2.234 | 0.148 | 13.401 | 0.300 | -1.463 | 1.202 | -15.635 | 0.346 |
| Sample 0079 | -1.743 | 0.161 | 13.100 | 0.199 | -1.944 | 1.076 | -14.843 | 0.270 |
| Sample 0080 | -1.841 | 0.171 | 13.180 | 0.208 | -2.161 | 1.063 | -15.022 | 0.276 |
| Sample 0081 | -1.746 | 0.111 | 13.133 | 0.147 | -1.339 | 1.042 | -14.879 | 0.193 |
| Sample 0082 | -1.445 | 0.186 | 13.298 | 0.174 | -3.207 | 0.713 | -14.743 | 0.223 |
| Sample 0083 | -2.800 | 0.172 | 13.061 | 0.414 | -2.102 | 1.343 | -15.861 | 0.480 |
| Sample 0084 | -2.665 | 0.187 | 13.223 | 0.414 | -1.935 | 1.346 | -15.888 | 0.471 |
| Sample 0085 | -2.906 | 0.248 | 13.193 | 0.470 | -2.134 | 1.396 | -16.099 | 0.556 |
| Sample 0086 | -2.731 | 0.196 | 13.029 | 0.426 | -2.121 | 1.401 | -15.759 | 0.498 |
| Sample 0087 | -1.601 | 0.200 | 13.072 | 0.214 | -2.286 | 1.097 | -14.672 | 0.285 |
| Sample 0088 | -1.656 | 0.168 | 13.036 | 0.188 | -2.107 | 1.066 | -14.691 | 0.253 |
| Sample 0089 | -1.698 | 0.139 | 13.389 | 0.165 | -1.930 | 1.002 | -15.087 | 0.222 |
| Sample 0090 | -1.871 | 0.163 | 13.297 | 0.218 | -1.584 | 1.148 | -15.168 | 0.288 |

Table S10: Summarized posteriors from the Beverton-Holt model fitting, showing means and standard deviations for the log-transformed paramaters $r_0$, $K$, $d$ and $\alpha$. Note the large standard deviations for the death rate ($d$) estimates.
